# Neural signals encoded by specific frequencies of closed-loop electrical stimulation structurally and functionally reconstruct spinal sensorimotor circuits after spinal cord injury

**DOI:** 10.1101/2022.05.10.491346

**Authors:** Kai Zhou, Dan Yang, Wei Wei, Hui Zhang, Wei Yang, Yunpeng Zhang, Yingnan Nie, Mingming Hao, Ting Zhang, Shouyan Wang, Yaobo Liu

## Abstract

Epidural electrical stimulation restores locomotion in animals and humans with spinal cord injury (SCI). However, the coding rules underlying electrical stimulation remain poorly understood, which has greatly limited the application of such electrical neuromodulation techniques in SCI clinical treatment. To elucidate the coding rules of electrical stimulation on spinal sensorimotor circuit reconstruction after complete SCI, we initially developed a spinal−muscle closed-loop stimulation protocol to mimic feedforward and feedback electrical signals in spinal sensorimotor circuits. Afterwards, using methods of sensorimotor function evaluation, neural circuit tracing and neural signal recording, we discovered a unique stimulus frequency of 10−20 Hz under closed-loop conditions was required for structural and functional reconstruction of spinal sensorimotor circuits. The single-cell transcriptome analysis of activated motoneurons characterized molecular networks involved in spinal sensorimotor circuit reconstruction. This study provides insights into neural signal decoding during spinal sensorimotor circuit reconstruction, and indicates a technological approach for the clinical treatment of SCI.

## Introduction

Electrical stimulation can augment or modify neuronal function and can have therapeutic benefits for certain neurological disorders ^1–4^. There is evidence that enhancing spinal excitability with either epidural or transcutaneous stimulation can restore some volitional motor output after spinal cord injury (SCI) ^5, 6^. Lumbosacral epidural stimulation can restore locomotor and autonomic function in both rodents and humans with SCI ^7^. When combined with overground locomotor training enabled by a weight-supporting device, epidural electrical stimulation (EES) promotes extensive reorganization of residual neural pathways that improves locomotion after stopping stimulation ^8–10^. Although it is exceedingly impressive that a technological platform was established to optimize neuromodulation in real time to achieve high-fidelity control of leg kinematics during locomotion in animals and patients, the parameters that influence the regenerative effects of electrical stimulation in peripheral and central regeneration are poorly understood. It is not clearly known (1) whether neural circuits can be reassembled constructively and functionally with closed-loop electrical stimulation (CLES) or with open-loop electrical stimulation (OLES), (2) whether and what characteristic of effective electrical stimulation are required for neural circuit reassembly, and (3) what molecular mechanisms are involved. These unanswered questions have greatly limited the application of electrical neuromodulation techniques in SCI clinical treatment.

To address these issues, we used the spinal sensorimotor circuits corresponding to the contraction of the hindlimb flexor, the tibialis anterior (TA) muscle, as the focus of our study. A CLES system was established using feedforward EES ^7, 11, 12^ combined with TA electrical stimulation ^13, 14^ feedback. Complete T9 spinal cord transection led to the complete interruption of supraspinal input to the lumbosacral neural circuit, providing an ideal research model to investigate the neuroplasticity of these spinal sensorimotor circuits under electrical stimulation. This study moves closer toward fully decoding the role of neuromodulation in neural circuit reassembly and indicates a potential framework for the clinical application of neural trauma treatment.

## Results

### Establishment of the closed-loop electrical stimulation (CLES) system

In mice with complete spinal cord injury (SCI), interruption of supraspinal innervation leads to disassembly of spinal sensorimotor circuits below the level of injury, which transmit neural signals between the spine and muscles and control the coordinated contraction of flexor and extensor muscles (**Figure 1A**) ^15^. Previous studies demonstrated the close-loop feedforward and feedback of electrical signals in spinal sensorimotor circuits is necessary to maintain its sensorimotor function output in intact mice ^16–18^. To investigate whether CLES leads to reassembly of spinal sensorimotor circuits, we selected TA sensorimotor reflex circuits corresponding to the contraction of the hindlimb flexor TA muscle as the focus of our study. We established a CLES system to mimic sensory feedback and feedforward muscle contraction loops with the integration of spinal (S1) and muscle (S2) stimulation, where the single S1 stimulation was defined as open-loop electrical stimulation-spinal cord stimulation (OLES-SCS) and a single S2 stimulation was defined as open-loop electrical stimulation-muscle stimulation (OLES-MS) (**Figure 1B**). To determine the proper implantation location of epidural electrodes in the spinal cord, retrograde, adeno-associated viral (AAV) vector carrying GFP (retro-AAV-GFP) was injected into the TA muscle of uninjured mice to retrogradely trace the corresponding motoneurons in the spinal cord. The motoneurons innervating the TA muscle were mainly distributed in the L2−L4 segment of the spinal cord based on analysis of continuous coronal and sagittal sections of the lumbar spinal cord (**Figure 1C, D**). A fine needle electrode was designed and manufactured to minimize damage to the mice caused by electrode transplantation. In mice with T9 spinal transection, this epidural electrode was placed in the dura mater of the L2−L4 segment of the spinal cord as S1 stimulation electrode during the T9 segment transection surgery to avoid secondary damage due to electrode implantation (**Figure 1B, E**). Magnetic resonance imaging (MRI) detection was used to determine the accuracy of the position of the implanted electrode in the spinal cord 4 weeks later (**Figure 1E**). A peripheral electrode was synchronously implanted only in the unilateral TA muscle of the hindlimb as S2 stimulation electrode (**Figure 1B**).

**Figure 1.**
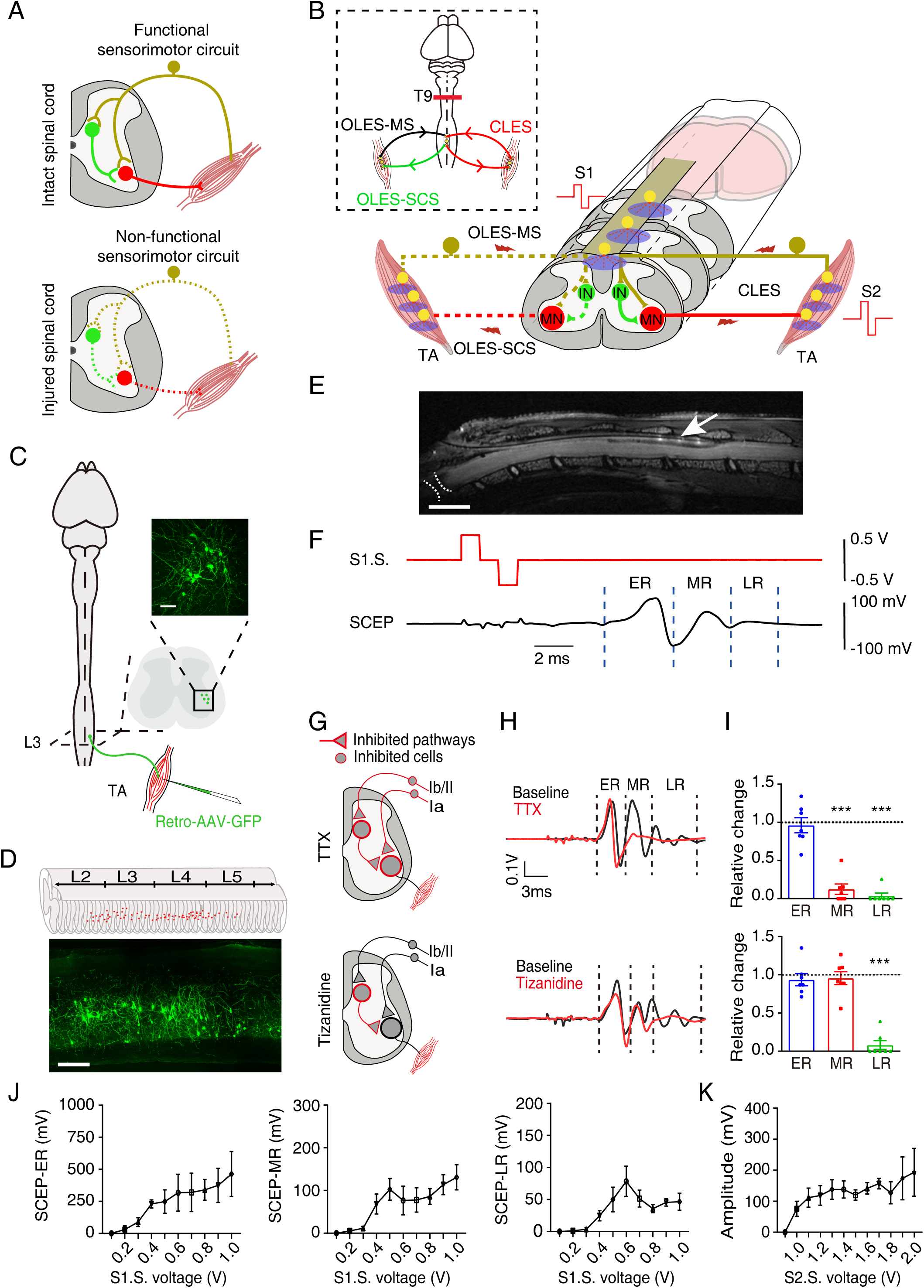
Characteristics of the CLES system. (**A**) Sensorimotor neural circuits in intact and SCI mice. (**B**) Schematic diagram showing OLES-SCS, OLES-MS, and CLES of the sensorimotor neural circuit in SCI mice. CLES system consists of two parts: S1, spinal cord epidural electrical stimulation; S2, muscle electrical stimulation. OLES-SCS consists only of S1; OLES-MS consists only of S2. The dashed box indicates that SCI mice were trained by CLES (red), OLES-SCS (green), and OLES-MS (black). IN, interneuron. MN, motoneuron. TA, tibialis anterior. (**C**) Retro-AAV-GFP was injected into TA muscles of intact mice to label motoneurons. Scale bar, 100 μm. (**D**) The spatial distribution of motoneurons innervating TA muscles. Scale bar, 200 μm. (**E**) The position of the electrode was confirmed by MRI analysis at 4 weeks after SCI; the white arrow points to sites of contact of the electrode contacts, and the dashed white lines indicate the SCI site. Scale bar, 2.5 mm. (**F**) S1 stimulation (red line) was administered, and TA muscles received SCEPs (black line) in intact mice. (**G**) Schematics illustrating which neurons, fibers, and/or circuits were likely inhibited in response to each experimental manipulation in anesthetized intact mice. (**H**) Representative SCEPs recorded in different mice during repeated S1 stimulation (1 Hz) and after the administration of TTX or tizanidine. Each waveform is the average of the responses to 50 stimuli. (**I**) Histogram plots for TTX (upper) and tizanidine (lower) conditions showing the relative change in the SCEPs as compared with baseline (n = 7 mice). (**J**) The SCEP curve for the ER, MR, and LR induced by S1 (n = 8 mice). (**K**) Curve showing the electrical signal received in the spinal cord by S2 (n = 5 mice). Data are shown as the mean ± SEM, \*\*\**p* < 0.001, statistical analysis was carried out with Student’s *t*-test.

To analyze the conduction characteristics of electrical signals in the TA sensorimotor reflex circuit under normal physiological conditions, S1 stimulation was administered to the dorsal spinal cord (L2−L4) along the midline in intact mice in an anesthetized state. A single pulse (0.2 ms) of S1 stimulation elicited spinal cord evoked potentials (SCEPs) in TA muscles; the recorded SCEPs had early latency responses (ERs), medium latency responses (MRs), and late latency responses (LRs) (**Figure 1F**). Pharmacological tests were subsequently conducted to isolate the ERs, MRs, and LRs corresponding to feedforward transmission in spinal sensorimotor circuits. ERs were not abolished by blocking sodium current through a local intrathecal injection of tetrodotoxin (TTX) at the L5/L6 spinal segments, but MRs and LRs were abolished (**Figure 1G-I**). Additionally, the amplitude of the LRs decreased substantially after an intrathecal injection of the high-dose α2-adrenergic receptor agonist tizanidine, but neither ERs nor MRs were affected (**Figure 1G-I**). These results indicated that only the LRs of SCEPs corresponded to the polysynaptic response in feedforward transmission mediated through Group II/Ib interneurons of spinal sensorimotor circuits. Consequently, the SCEPs were analyzed at disparate S1 stimulation intensities to determine the stimulus intensity of S1 (**Figure 1J**). LR amplitudes reached a peak when the S1 stimulation intensity was 0.6 V, indicating that feedforward transmission mediated by spinal sensorimotor circuits was fully activated. Considering the individual differences among mice, S1 stimulation intensity was adjusted between 0.5 V and 0.7 V in the CLES system to transmit feedforward signals to the muscle. Spinal epidural recording confirmed the effective response (unimodal peak) with minimal stimulation of 1 V administered to the TA muscle (**Figure 1K**). Thus, TA muscles received 1 V of stimulation in the CLES system to transmit sensory feedback signals to the spinal cord.

### Restoration of polysynaptic electrical conduction in spinal sensorimotor circuits by CLES in SCI mice

Previous studies have shown that the frequency of electrical stimulation is one of the key parameters to determine the effect of electrical neuromodulation, and it is found low-frequency electrical stimulation promotes the recovery of sensorimotor function after SCI ^19–21^. Therefore, we chose to identify the frequency characteristics of effective electrical stimulation in the frequency range of 1-40 Hz. Electrical stimulation training started one week after SCI and continued for three weeks (**Figure 2A**). SCEPs were detected in the TA muscle (**Figure 2A**). The results show that MRs and LRs were clearly observed in the sham group and had almost disappeared in the untrained group, although ERs showed no changes (**Figure 2B-J**). This suggests that polysynaptic electrical conduction in the sensorimotor circuits was interrupted by SCI. MRs and LRs were almost undetectable in the 1- to 40-Hz OLES-SCS, and 1- to 40-Hz OLES-MS groups 4 weeks after SCI (**Figure 2B-G**). However, in the 10- to 20-Hz CLES groups, MR and LR amplitudes increased significantly after 3 weeks of CLES training as compared with mice in the untrained group (**Figure 2H-J**).

**Figure 2.**
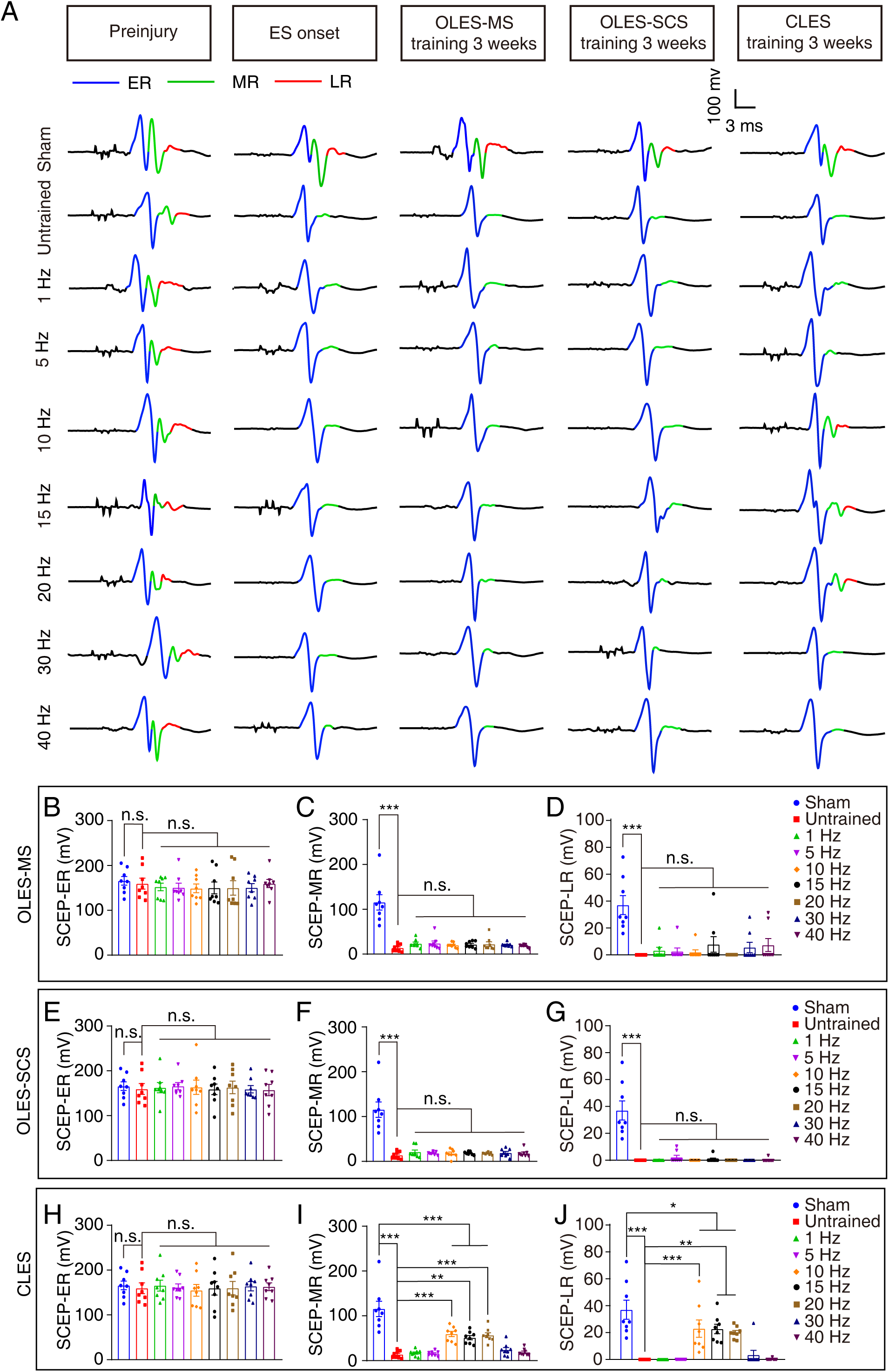
SCEPs recorded in the TA muscle of SCI mice after CLES. (**A**) SCEPs were recorded in sham, untrained, and after 3 weeks of 1- to 40-Hz CLES, 1- to 40-Hz OLES-SCS, and 1- to 40-Hz OLES-MS training. (**B**-**D**) The amplitudes of ERs (**B**), MRs (**C**), and LRs (**D**) of SCEPs in the 1- to 40-Hz OLES-MS, untrained, and sham groups (n = 8 mice). (**E**-**G**) Histogram reporting the amplitudes of ERs (**E**), MRs (**F**), and LRs (**G**) of SCEPs in the 1- to 40-Hz OLES-SCS, untrained, and sham groups (n = 8 mice). (**H**-**J**) Histogram reporting the amplitudes of ERs (**H**), MRs (**I**), and LRs (**J**) of SCEPs in the 1- to 40-Hz CLES, untrained, and sham groups (n = 8 mice). Data are shown as the mean ± SEM, ns: no statistical difference, \**p* < 0.05, \*\**p* < 0.01, \*\*\**p* < 0.001, one-way ANOVA followed by Bonferroni’s post-test.

To clarify the effect of spinal cord high-voltage stimulation training, 2 V high-voltage electrical stimulation was applied to mice in the OLES-SCS group (OLES-SCShv), which was the maximum limit that mice can tolerate (data not shown). The results showed that MR and LR amplitudes were not significantly improved in the 1- to 40-Hz OLES-SCShv groups compared with the untrained group at 4 weeks after SCI (**Figure 2-figure supplement 1A-F**). In summary, the results show that polysynaptic electrical conduction in sensorimotor circuits was restored by only 10- to 20-Hz CLES. 10- to 20-Hz CLES were identified as effective electrical stimulations, whereas 1-Hz, 5-Hz, 30-Hz, and 40-Hz CLES were identified as ineffective electrical stimulations.

### Restoration of sensorimotor function of hind limbs in SCI mice by 10- to 20-Hz CLES

To further test the effect of CLES, OLES-SCS, OLES-MS and OLES-SCShv on sensorimotor function in SCI mice, a basso mouse scale (BMS) score and Von Frey test were performed under different experimental conditions. BMS test was performed to evaluate the overall basic locomotor performance. Mice were observed in an open field and subjected to BMS scoring at 0, 1, 3, 7, 14, 21, and 28 days after injury. Compared with that of the untrained group, the motor function of the hind limbs was significantly restored by 10- to 20-Hz CLES (**Figure 3A**). In contrast, mice in the 1- to 40-Hz OLES-SCS, 1- to 40-Hz OLES-MS, and 1- to 40-Hz OLES-SCShv groups showed no significant improvement in hind limb locomotor function as compared with mice in the untrained group (**Figure 3B, C and Figure 3-figure supplement 1A**). Next, motor function of the hind limbs was evaluated in the 10- to 20-Hz OLES-SCS, 10- to 20-Hz OLES-MS, 10- to 20-Hz OLES-SCShv, and 10- to 20-Hz CLES groups with BMS scoring. Compared with the 10- to 20-Hz OLES−SCS, 10- to 20-Hz OLES−MS, and 10- to 20-Hz OLES-SCShv groups, motor function recovery of the hind limbs was increased by 10- to 20-Hz CLES (**Figure 3D**). But under other frequencies of electrical stimulation, compared with the OLES-SCS, OLES-MS, and OLES-SCShv groups, the motor function of the hind limbs of mice in the CLES group was not significantly increased (**Figure 3-figure supplement 2A-D**). Thus, these results indicate that the closed loop was necessary for the improvement of locomotor function after SCI, and functional motor recovery was effectively promoted by 10- to 20-Hz CLES instead of OLES after SCI.

**Figure 3.**
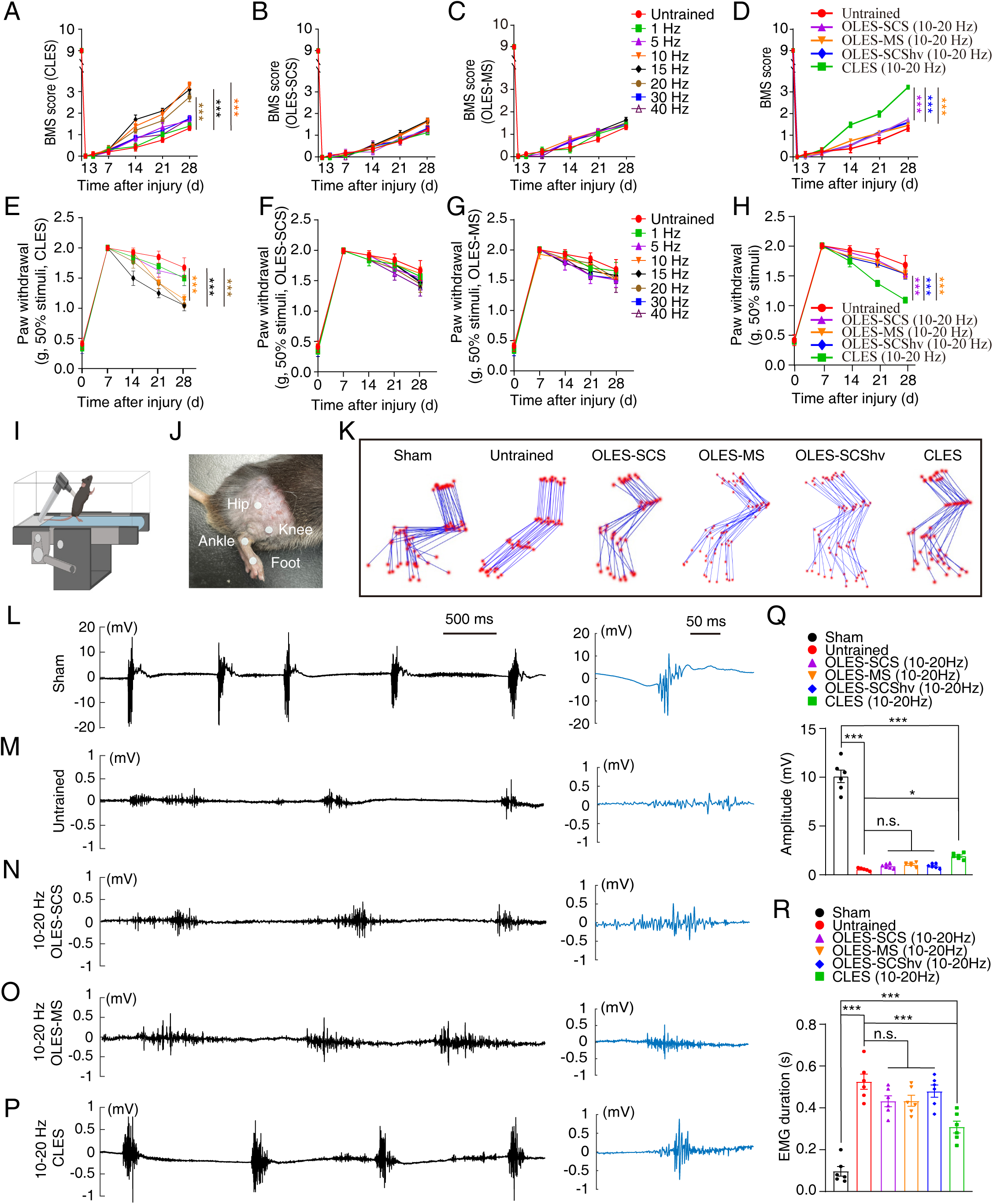
Detection of sensory and motor function of hind limbs in SCI mice after CLES and OLES. (**A**-**C**) Hindlimb BMS scores of mice in the untrained, 1- to 40-Hz CLES (**A**), 1- to 40-Hz OLES-SCS (**B**), and 1- to 40-Hz OLES-MS (**C**) groups at different time points after SCI (n = 8 mice). (**D**) Weighted average comparisons of the hindlimb BMS scores of mice in the untrained, 10−20 Hz OLES-SCS, 10−20 Hz OLES- MS, 10−20 Hz OLES-SCShv, and 10−20 Hz CLES groups at different time points after SCI (n = 9 mice, there were three mice in each frequency group). (**E**-**G**) The mechanical pain threshold of hind limbs in the untrained, 1- to 40-Hz CLES (**E**), 1- to 40-Hz OLES- SCS (**F**), and 1- to 40-Hz OLES-MS (**G**) groups at different time points after SCI (n = 8 mice). (**H**) The line graph reports the weighted average comparison of the mechanical pain threshold of the hind limbs in the untrained, 10−20 Hz OLES-SCS, 10−20 Hz OLES-MS, 10−20 Hz OLES-SCShv, and 10−20 Hz CLES groups at different time points after SCI (n = 9 mice, there were three mice in each frequency group). (**I**) While mice moved on the treadmill, EMG signals from the TA muscle were recorded. (**J**) The hip, knee joint, ankle joint, and sole of the foot of each mouse hindlimb was marked. (**K**) Representative stick diagram decomposition of hindlimb movements in the sham, untrained, 10−20 Hz OLES-SCS, 10−20 Hz OLES-MS, 10−20 Hz OLES-SCShv, and 10−20 Hz OLES groups, together with 15-frame successive trajectories of the limb endpoint. (**L-P**) Left, the graphs show surface EMGs for TA muscles in the sham (**L**), untrained (**M**), 10−20 Hz OLES-SCS (**N**), 10−20 Hz OLES-MS (**O**), and 10−20 Hz CLES (**P**) groups within 5s. Right, the representative waveform on the left. (**Q**) The maximum amplitude of surface EMG bursts continuously for 5 seconds in the sham, untrained, 10−20 Hz OLES-SCS, 10−20 Hz OLES-MS, 10−20 Hz OLES-SCShv, and 10−20 Hz CLES groups (n = 6 mice, there were two mice in each frequency group). (**R**) Average duration of surface EMG bursts in 5 seconds in the sham, untrained, 10−20 Hz OLES-SCS, 10−20 Hz OLES-MS, 10−20 Hz OLES-SCShv, and 10−20 Hz CLES groups (n = 6 mice, there were two mice in each frequency group). Data are shown as the mean ± SEM, ns: no statistical difference, \**p* < 0.05, \*\*\**p* < 0.001, two-way ANOVA followed by Fisher’s least significant differences post-test (LSD) (**A-H**), one-way ANOVA followed by Bonferroni’s post-test (**Q** and **R)**.

The response elicited by the application of von Frey filaments to the skin can be used as a measure of tactile sensitivity. Mice were observed in a transparent box, and von Frey filaments were applied with different forces to stimulate the paws of the hind limbs at 0, 7, 14, 21, and 28 days after injury. The mechanical pain threshold of the hind limbs of mice increased significantly after SCI (**Figure 3E-G**). In contrast, the mechanical pain threshold of the hind limbs in the 10- to 20-Hz CLES mice was significantly reduced after the 3-week training period (**Figure 3E**). Mice in the 1- to 40-Hz OLES-SCS, 1- to 40-Hz OLES-MS, and 1- to 40-Hz OLES-SCShv groups showed no significant difference in the mechanical pain threshold of the hind limbs as compared with that of mice in the untrained group (**Figure 3F, G and Figure 3-figure supplement 1B**). The mechanical pain threshold of the hind limbs was significantly reduced in the 10- to 20-Hz CLES groups as compared with that of the hind limbs in the 10- to 20-Hz OLES-SCS, 10- to 20-Hz OLES-MS, and 10- to 20-Hz OLES-SCShv groups (**Figure 3H**). But under other frequencies of electrical stimulation, compared with the OLES-SCS, OLES-MS, and OLES-SCShv groups, the mechanical pain threshold of the hind limbs of mice in the CLES group was not significantly decreased (**Figure 3-figure supplement 2E-H**). These results indicate that the sensory function of the hind limbs was restored by 10- to 20-Hz CLES after SCI.

To detect electromyography (EMG) bursts during rhythmic movement of mice, we designed a body weight−supporting (BWS) device combined with a treadmill so that the hind limbs of the mice just touched the track of the treadmill. The treadmill operated at a speed of 2 m·minute^−1^, and EMG signals from the TA muscles were observed when the mice were moving (**Figure 3I**). For these experiments, we marked the hip, knee, ankle joint, and sole of the foot of each mouse hind limb to determine movement trajectories (**Figure 3J**). In mice of the untrained group, the ankle joint of the hind limb was barely flexed (**Figure 3K**). However, the extent of dorsiflexion of the hind limb ankle joint was greater in the 10- to 20-Hz CLES groups relative to that in the 10- to 20-Hz OLES groups and was more similar to that of the sham group (**Figure 3K**). Next, we recorded the EMG bursts of the mice during movement (**Figure 3L-P**). With the sham group mice serving as a control (**Figure 3L and Figure 3-figure supplement 1C**), we could hardly detect EMG bursts in the TA muscle of the untrained, 1- to 40-Hz OLES-SCS, 1- to 40-Hz OLES-MS, and 1- to 40-Hz OLES-SCShv groups mice (**Figure 3M-O and Figure 3-figure supplement 1D, E**). The amplitude of TA EMG bursts in the 10- to 20-Hz CLES groups was significantly higher than that of the untrained group (**Figure 3Q**). The duration of a single EMG was also measured and was significantly longer in the untrained group than that in the sham group (**Figure 3R**). However, the duration of a single EMG recorded in the 10- to 20-Hz CLES groups was significantly shorter than that in the untrained group (**Figure 3R**). Together, 10- to 20- Hz CLES enhanced the electrical signal emission of spinal cord neurons to peripheral muscles after SCI.

To accurately and objectively evaluate the effect of CLES on TA muscle contraction function after SCI, a newly developed pressure sensor that does not affect locomotion in intact mice was used to measure the intensity and frequency of the TA muscle contraction. In the sham, untrained, 1- to 40-Hz OLES-SCS, 1- to 40-Hz OLES- MS and 1- to 40-Hz OLES-SCShv groups, the contraction function of the TA muscle was detected by the pressure sensor (**Figure 4A-I and Figure 4-figure supplement 1A-I**). SCI resulted in a significant reduction in the intensity and frequency of the TA muscle contraction relative to those parameters in sham group mice (**Figure 4J-O and Figure 4-figure supplement 1J, K**). As compared with the untrained group, both the frequency and the intensity of activity significantly increased in the 10- to 20-Hz CLES groups (**Figure 4J, K**). In contrast, TA muscle contraction function was not significantly improved in the 1- to 40-Hz OLES-SCS, 1- to 40-Hz OLES-MS, and 1- to 40-Hz OLES-SCShv groups as compared with mice in the untrained group (**Figure 4L-O and Figure 4-figure supplement 1J, K**). These results indicate that TA muscle contractility function was effectively recovered by 10- to 20-Hz CLES, but not by OLES after SCI.

**Figure 4.**
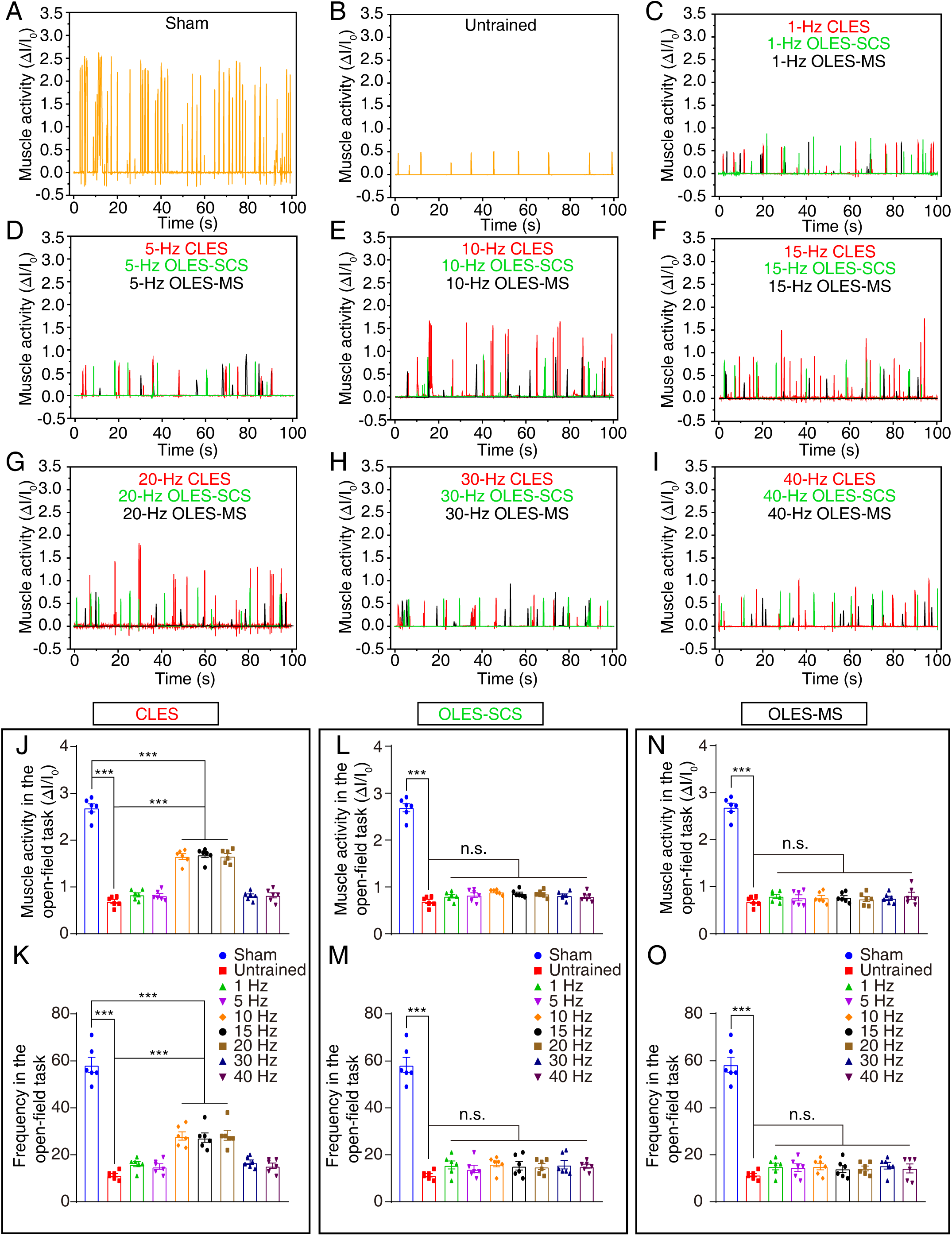
Evaluation of TA muscle contraction function in SCI mice after CLES and OLES. (**A** and **B**) TA muscle contraction and its corresponding relative current in mice in the sham (**A**), untrained (**B**). (**C-I**) TA muscle contraction curve of mice in 1- to 40- Hz CLES (red), OLES-SCS (green) and OLES-MS (black) groups. (**J** and **K**) The intensity (**J**) and frequency (**K**) of TA muscle contraction in mice in the sham, untrained, and 1- to 40-Hz CLES groups (n = 6 mice). (**L** and **M**) The intensity (**L**) and frequency (**M**) of TA muscle contraction in mice in the sham, untrained, and 1- to 40- Hz OLES-SCS groups during the open-field task (n = 6 mice). (**N** and **O**) The intensity (**N**) and frequency (**O**) of TA muscle contraction in mice in the sham, untrained, and 1- to 40-Hz OLES-MS groups (n = 6 mice). Data are shown as the mean ± SEM, ns: no statistical difference, \*\*\**p* < 0.001, one-way ANOVA followed by Bonferroni’s post-test.

In summary, in vivo functional analyses provide evidences that indicate the sensorimotor function of the hind limbs was restored through 10- to 20-Hz CLES instead of OLES after SCI.

### Reconstruction of hind limb neuromuscular junctions (NMJs) in SCI mice by 10- to 20-Hz CLES

To verify that the spinal sensorimotor circuit had been reconstructed as a result of CLES, we conducted a separate examination of each component of the circuit. We first observed the TA muscle fibers and NMJs formed between the motor nerve and TA muscle (**Figure 5A**). NMJs were labeled by immunofluorescence staining with antibodies that recognized. The coronal section of the TA muscles was stained with α-bungarotoxin (α-BTX, red) to label AChR and anti-neurofilament/synapsin (NF/Syn, green/blue) antibodies to label axon terminals (**Figure 5B and Figure 5-figure supplement 1**). Hematoxylin and eosin (H&E) staining analysis of muscle tissue was performed to evaluate the degree of muscle atrophy after SCI (**Figure 5B and Figure 5-figure supplement 1**). Compared to sham group, the cross-sectional area of muscle fibers decreased significantly in the untrained and CLES onset groups (**Figure 5C**). In addition, compared to CLES onset group, the cross-sectional area of the muscle fiber decreased significantly in the untrained group, indicating persistent muscle atrophy four weeks after SCI (**Figure 5C**). The cross-sectional area of the muscle fibers increased significantly in 10- to 20-Hz CLES groups compared to untrained group, indicating that 10- to 20-Hz CLES effectively rescued muscular atrophy after SCI (**Figure 5C**). Next, we compared the NMJs of different CLES groups. We randomly selected six TA muscle endplates in the field of view, and calculated the percentage of fully innervated, partially innervated, and denervated endplates (fully innervated, syn and α-BTX overlap; partial innervation, syn and α-BTX partially overlap; denervated, syn and α-BTX do not overlap). There were fewer endplates in the TA muscle that was fully innervated in the untrained (10.2%) and CLES onset (20%) group than in the sham group (92.7%) (**Figure 5D**). Notably, fully innervated endplates were increased in the 10- to 20-Hz CLES groups (46.8%, 56.6%, 43.2%) (Fig. 5d). Together, these results suggest that NMJ deficiency in SCI mice was effectively restored and muscle atrophy was reduced by 10- to 20-Hz CLES.

**Figure 5.**
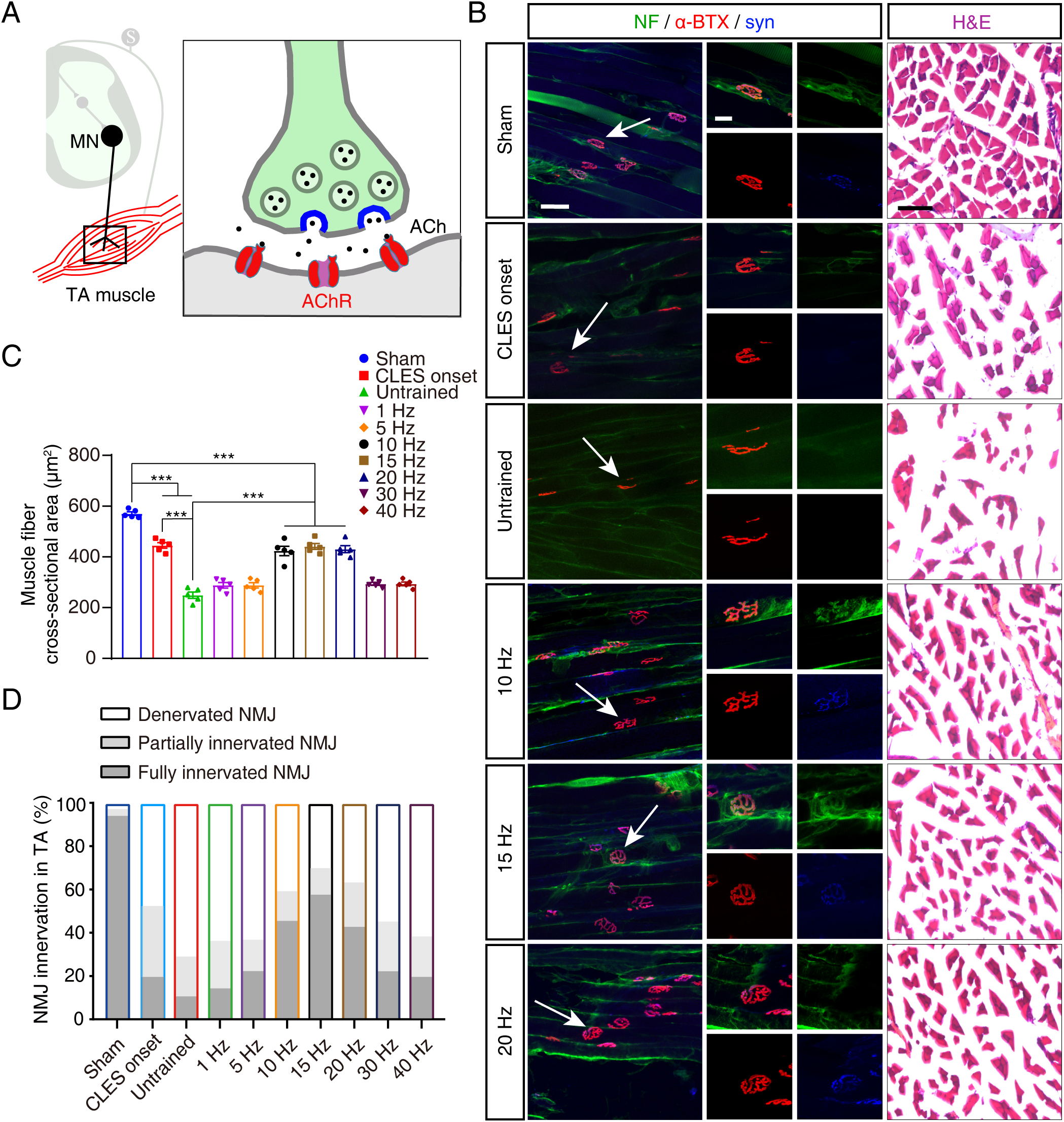
The morphology and neural innervation of TA muscles in SCI mice after CLES. (**A**) Schematic diagram of the NMJ. right, Green: motoneuron axons; blue: presynaptic synaptic vesicle protein; red: acetylcholinergic receptor. (**B**) Left, antibodies against NF (green), α-BTX (red), and syn (blue) were used to label NMJs on the TA muscle of hind limbs from mice in the sham, CLES onset, untrained, and 10- to 20-Hz CLES groups. The white arrow indicates a single NMJ in each image. Scale bar, 50 μm. Middle, a higher-magnification image of the NMJ indicated by the white arrow. Scale bar, 20 μm. Right, H&E staining of the TA muscle of hind limbs from mice in the sham, CLES onset, untrained, and 10- to 20-Hz CLES groups. Scale bar, 50 μm. (**C**) The cross-sectional area of muscle fibers in the TA muscle of hind limbs from mice in the sham, CLES onset, untrained, and 1- to 40-Hz CLES groups (n = 5 mice). (**D**) The percentage of denervation, partial innervation, and complete innervation of the TA muscles of hind limbs from mice in the sham, CLES onset, untrained, and 1- to 40-Hz CLES groups (n = 5 mice). Data are shown as the mean ± SEM, \*\*\**p* < 0.001, one-way ANOVA followed by Bonferroni’s post-test (**c**).

### Reconnection of hind limb spinal sensory-motor connectivity in SCI mice by 10- to 20-Hz CLES

Subsequently, we observed the projection of sensory neuron axons to the spinal cord. Cholera toxin B (CTB) tracer was injected into the TA muscles of SCI mice, and these mice were euthanized 2 weeks later to trace the proprioceptive axon terminals ^22^ (PTs) in the spinal cord (**Figure 6A**). Three-dimensional reconstruction was performed to analyze the labeled PTs in the spinal cord (**Figure 6B and Figure 6-figure supplement 1**). Compared to sham group, a significant reduction in PTs was observed in the untrained and CLES onset groups, indicating that the projection of sensory neurons to the spinal cord was disrupted one week after SCI (**Figure 6C**). In contrast, PTs were restored significantly in the 10- to 20-Hz CLES groups compared to untrained group (**Figure 6C**). These results suggest that 10- to 20-Hz CLES effectively promoted the innervation of proprioceptive axons in the spinal cord after SCI.

**Figure 6.**
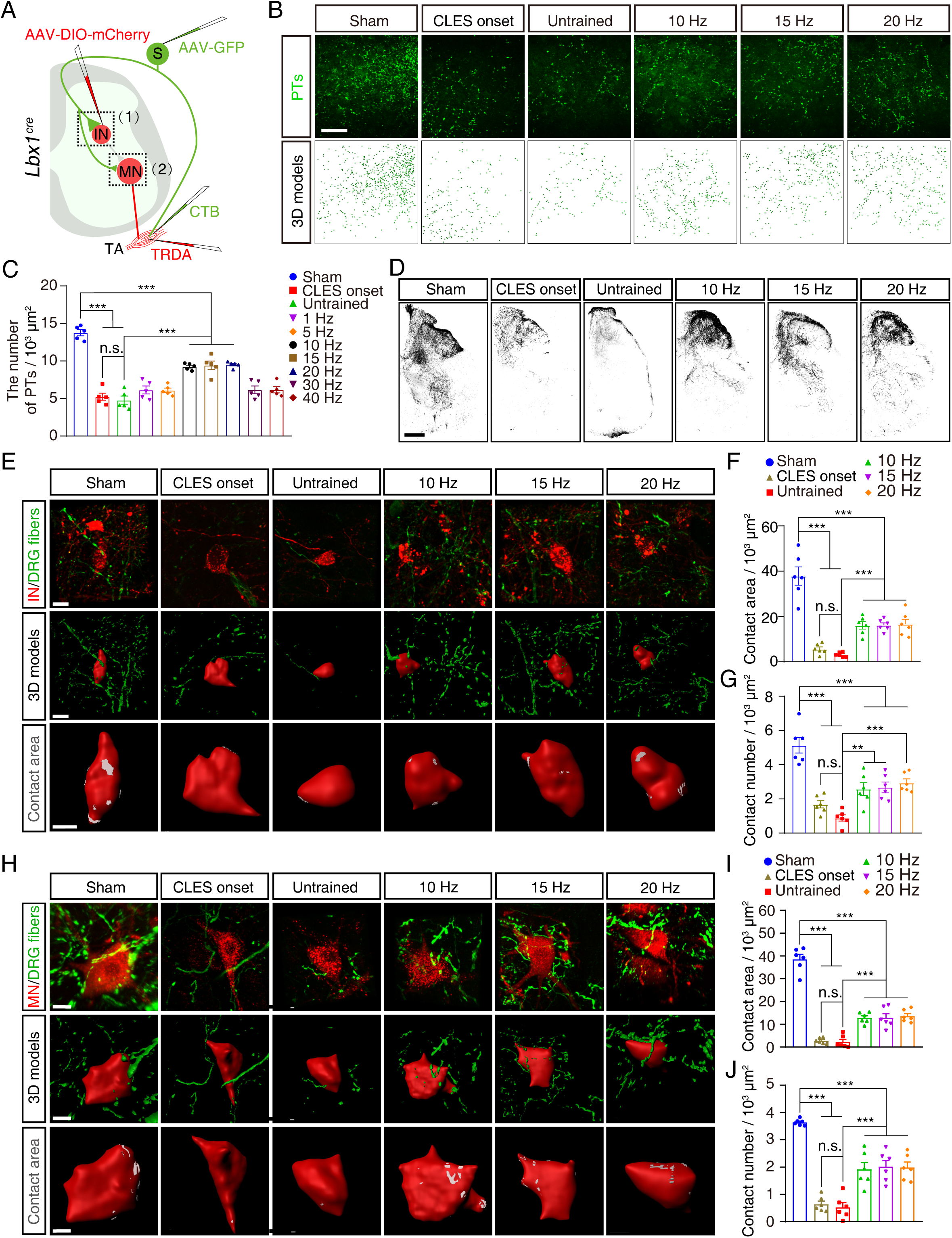
Sensory-motor connectivity in SCI mice after CLES. (**A**) The diagram illustrates the tracing of PTs (1), interneurons (1), motoneurons (2), and DRG fibers (1 and 2) in *Lbx1^cre^* mice. MN, motoneuron; S, sensory neuron; IN, interneuron. (**B**) CTB was injected into the TA muscle of mice in the sham, CLES onset, untrained, and 10- to 20-Hz CLES groups to anterogradely trace PTs (green) in the spinal cord (upper images), and PTs were subjected to three-dimensional (3D) modeling (lower images). Scale bar, 50 μm. (**C**) The number of PTs per 1000 μm^2^ of the coronal area of the spinal cord in the sham, CLES onset, untrained, and 1- to 40-Hz CLES groups (n = 5 mice). (**D**) AAV-GFP was injected into the DRG of mice in the sham, CLES onset, untrained, and 10- to 20-Hz CLES groups to anterogradely label the DRG fibers in the spinal cord. Scale bar, 200 μm. (**E**) Upper, colocalization of DRG fibers and interneurons in the sham, CLES onset, untrained, and 10- to 20-Hz CLES groups. Scale bar, 15 μm. Middle, three-dimensional reconstruction of DRG fibers and interneurons. Scale bar, 15 μm. Lower, highly magnified image of interneurons. The gray region represents the region of connection between DRG fibers and interneurons. Scale bar, 10 μm. (**F** and **G**) The contact area (**F**) and number (**G**) of contacts of DRG fibers per 1000 μm^2^ of interneuron surface area in the sham, CLES onset, untrained, and 10- to 20-Hz CLES groups (n = 6 mice). (**H**) Upper, colocalization of DRG fibers and motoneurons in the sham, CLES onset, untrained, and 10- to 20-Hz CLES groups. Scale bar, 15 μm. Middle, three- dimensional reconstruction of DRG fibers and motoneurons. Scale bar, 15 μm. Lower, highly magnified image of motoneurons. The gray region represents the region of connection between DRG fibers and motoneurons. Scale bar, 10 μm. (**I** and **J**) The contact area (**I**) and number (**J**) of contacts of DRG fibers per 1000 μm^2^ of motoneuron surface area in the sham, CLES onset, untrained, and 10- to 20-Hz CLES groups (n = 6 mice). Data are shown as the mean ± SEM, ns: no statistical difference, \*\**p* < 0.01, \*\*\**p* < 0.001, one-way ANOVA followed by Bonferroni’s post-test.

We next explored the specificity of monosynaptic sensory inputs onto motoneurons supplying TA muscles. We tracked the central projection of sensory neuron afferents by injecting AAV-GFP into the dorsal root ganglion (DRG) at the L2−L4 segment in mice (**Figure 6A**). In the sham group, DRG fibers labeled from the L2−L4 DRG were mainly distributed in the dorsal spinal cord and extended to the ventral spinal cord, where motoneurons reside (**Figure 6D**). In contrast, DRG fibers terminated in the intermediate spinal cord in untrained mice and mice at CLES onset (**Figure 6D**). Notably, DRG fibers were found to re-extend to the ventral spinal cord in the 10- to 20-Hz CLES groups (**Figure 6D**). These results indicate that the projection of sensory neuron axons to the ventral spinal cord was restored by 10- to 20-Hz CLES.

AAV-DIO-mCherry was injected into the intermediate area of gray matter at the L2−L4 segment of *Lbx1^cre^ mice* to label the interneurons (**Figure 6A**). We colocalized DRG fibers and interneurons and performed three-dimensional modeling (**Figure 6E**). The area and number of DRG fibers connected with interneurons were measured (**Figure 6E**). Compared with the sham group, the area and the number of connections between sensory afferent fibers and interneurons were significantly reduced in untrained and the CLES onset group mice, indicating the sensory-interneuron connections were destroyed one week after SCI (**Figure 6F, G**). As compared with the untrained group, the area and number of DRG fibers and interneuron connections increased significantly in the 10- to 20-Hz CLES groups (**Figure 6F, G**). These results indicate that the sensory-interneuron connections were rebuilt by 10- to 20-Hz CLES.

Texas Red−dextran amine (TRDA) was injected into the TA for retrograde tracing of motoneurons (**Figure 6A**). We colocalized DRG fibers and motoneurons and performed three-dimensional modeling (**Figure 6H**). The area and number of DRG fibers connected with motoneurons were measured (**Figure 6H**). Compared with the sham group, the area and the number of connections between sensory afferent fibers and motoneurons were significantly reduced in untrained and the CLES onset group mice, indicating the sensory-motor connections were destroyed one week after SCI (**Figure 6I, J**). As compared with the untrained group, the area and number of DRG fibers and motoneuron connections increased significantly in the 10- to 20-Hz CLES groups (**Figure 6I, J**). These results indicate that the sensory-motor connections were reconstructed by 10- to 20-Hz CLES.

### Morphological characterization of motoneurons restored by 10- to 20-Hz CLES

Retro-AAV-GFP was injected into the TA muscle of mice to retrogradely trace the corresponding motoneurons in the spinal cord (**Figure 7A and Figure 7-figure supplement 1A**). Compared to sham group, obvious loss of spinal motoneurons was observed in the untrained group, but not in the onset group (**Figure 7B**). In contrast, the most significant survival of motoneurons was observed in the 10- to 20-Hz CLES groups (**Figure 7B**). Three-dimensional reconstruction was applied to analyze the distal dendrites and somas of labeled motoneurons (**Figure 7A and Figure 7-figure supplement 1A**). The volume of motoneurons was reduced significantly in the untrained and CLES onset group, indicating that SCI led to motoneuron atrophy (**Figure 7C, D**). In contrast, motoneuron atrophy was effectively reversed in the 10- to 20-Hz CLES groups (**Figure 7C, D**). And, the volume of somas and dendrites of motoneurons in the 10- to 20- Hz CLES groups almost recovered to the preinjury level (**Figure 7C, D**). These results demonstrate that the application of 10- to 20-Hz CLES effectively prevented neuronal apoptosis and loss, and regenerated the branching structures of motoneurons after SCI.

**Figure 7.**
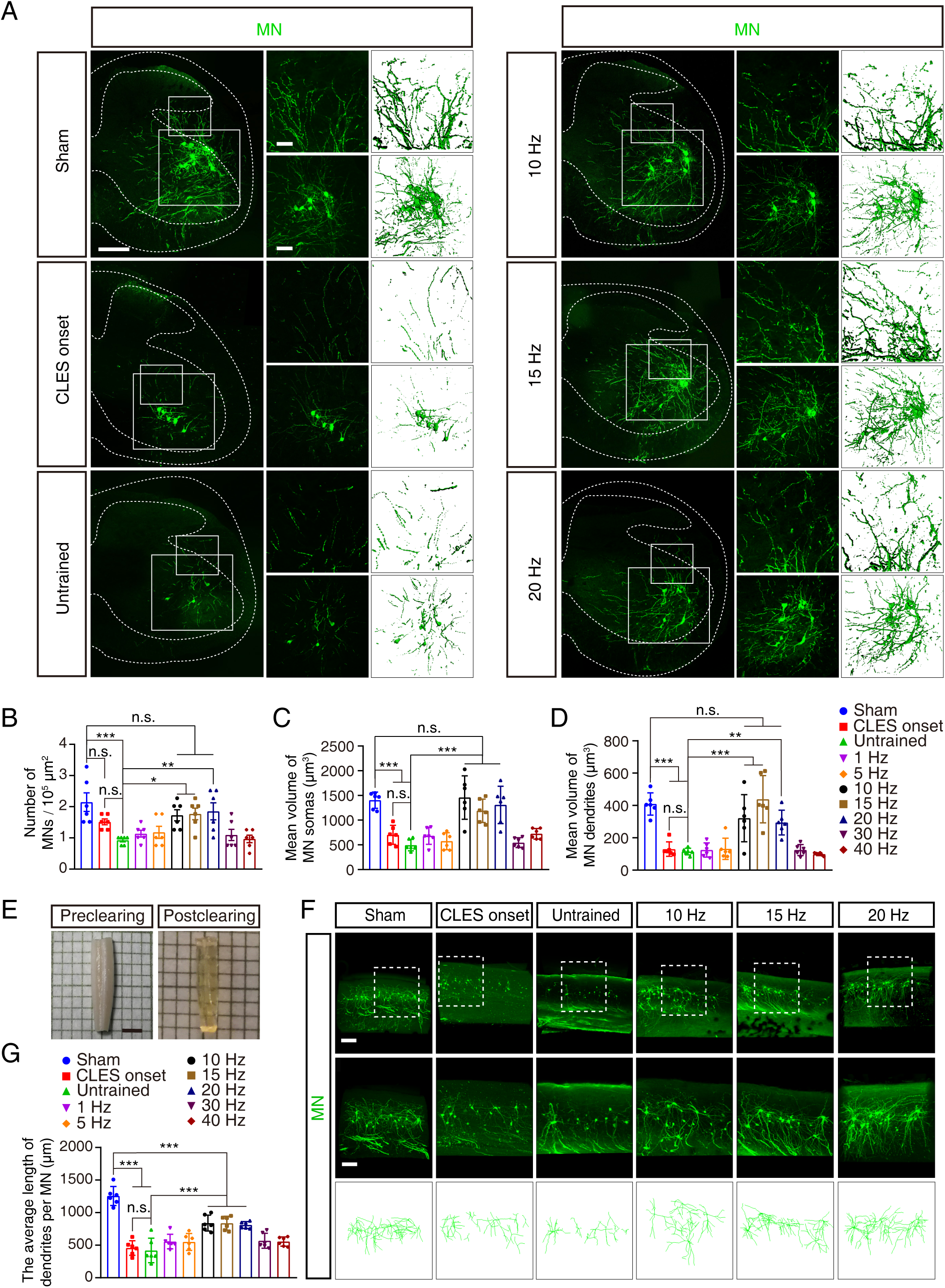
Morphological characteristics of spinal motoneurons in SCI mice after CLES. (**A**) Retro-AAV-GFP was injected into the TA muscle to retrogradely trace spinal motoneurons. Left, spinal motoneurons in the sham, CLES onset, untrained, and 10- to 20-Hz CLES groups. Scale bar, 200 μm. Upper right, higher-magnification image and three-dimensional model of the dendrites in the small boxed area. Scale bar, 50 μm. Lower right, the partial image and three-dimensional model of the motoneurons in the large boxed area. Scale bar, 100 μm. (**B**) The number of MNs per 100000 μm^2^ of the coronal area of the spinal cord in the sham, CLES onset, untrained, and 1- to 40-Hz CLES groups (n = 6 mice). (**C** and **D**) The mean volume of motoneuron somas (**C**) and dendrites (**D**) in the sham, CLES onset, untrained, and 1- to 40-Hz CLES groups (n = 6 mice). (**E**) The tissue clearing technique was performed on the lumbar spinal cord. Scale bar, 2 mm. (**F**) Upper, spatial distribution of motoneurons in L2−L5 spinal cord in the sham, CLES onset, untrained, and 10- to 20-Hz CLES groups. Scale bar, 200 μm. Lower, higher-magnification image and three-dimensional models of the boxed area. Scale bar, 100 μm. (**G**) The average length of dendrites per motoneuron in L2−L5 spinal cord in the sham, CLES onset, untrained, and 10- to 20-Hz CLES groups (n = 6 mice). Data are shown as the mean ± SEM, ns: no statistical difference, \**p* < 0.05, \*\**p* < 0.01, \*\*\**p* < 0.001, one-way ANOVA followed by Bonferroni’s post-test.

To further confirm the reassembly of the spinal sensorimotor circuit with 10- to 20-Hz CLES after SCI, retro-AAV-GFP was injected into the TA muscle to retrogradely trace the corresponding spinal motoneurons in L2−L5 spinal cord. At 2 weeks after the injection, the L2−L5 spinal cord was collected and made transparent by using a tissue optical clearing technique ^23^, and whole-mount immunostaining was carried out to visualize the labeled motoneurons and their branching structures (**Figure 7E, F and Figure 7-figure supplement 1B**). Three-dimensional reconstruction of the whole-mount staining images was conducted to comprehensively observe motoneuron dendrites and quantify their complexity (**Figure 7F and Figure 7-figure supplement 1B**). Compared to sham group, the average length of dendrites per motoneuron was significantly reduced in the untrained and CLES onset group, indicating that the complexity of motoneuron branching structures and the possibility of making connections with interneurons was reduced one week after SCI (**Figure 7G**). The average length of dendrites per motoneuron was significantly increased relative to the untrained mice in the 10- to 20-Hz CLES groups (**Figure 7G**). These results indicate that the application of 10- to 20-Hz CLES significantly promoted the complexity of motoneuron dendrites and increased the chance of synaptic reformation between interneurons and motoneurons after SCI.

### Restoration of premotor interneuron input into motoneurons in SCI mice by 10- to 20-Hz CLES

Premotor interneurons responsible for relaying signals are essential for the movement of mouse hindlimbs ^24^. Therefore, we were curious whether CLES affects the communication between premotor interneurons and motoneurons in SCI mice. Anterograde, AAV vector carrying rabies virus glycoprotein (RVG) and DIO promoter (AAV-DIO-RVG) was injected into the ventral spinal cord of *ChAT^cre^ mice*, RVG was expressed in motoneurons and enhanced green fluorescent protein−labeled and G-deleted rabies virus (RV-N2C (G)-ΔG-EGFP) was injected into the TA muscle to retrogradely trace the corresponding premotor interneurons (**Figure 8A**). Retro-AAV-mCherry was injected into the TA muscle to retrogradely trace motoneurons (**Figure 8A**). We found that the connections between premotor interneurons and motoneurons were distributed mainly in lamina VII of the Rexed’s laminae (**Figure 8B**). Axons of premotor interneurons and the dendrites of motoneurons were colocalized in lamina VII, and three-dimensional modeling was performed (**Figure 8C**). The area and number of synaptic connections between premotor interneurons and motoneurons were significantly reduced in the untrained and CLES onset group as compared with the sham group (**Figure 8D, E**). Compared to untrained group, the number and area of synaptic connections between premotor interneurons and motoneurons were significantly increased in the 10- to 20-Hz CLES groups (**Figure 8D, E**). These results indicate that 10- to 20-Hz CLES effectively promoted the reinnervation of premotor interneurons to motoneurons after SCI.

**Figure 8.**
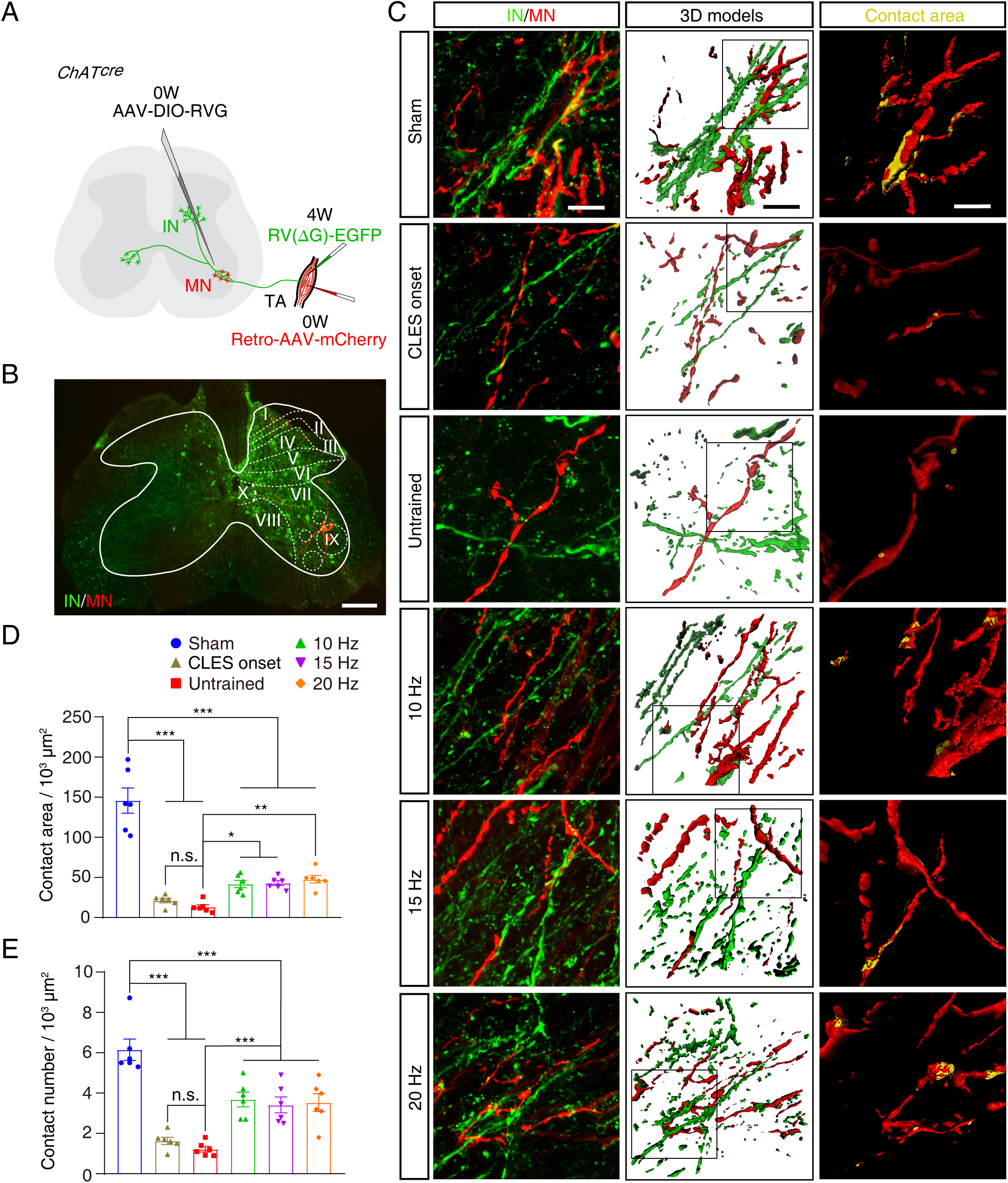
The connection between spinal interneurons and motoneurons in SCI mice after CLES. (**A**) Injection scheme to visualize the spinal interneurons network innervating the TA muscle using *ChAT^cre^* mice. (**B**) Interneurons (green) and motoneurons (red) were shown from an intact mouse. Spatial organization of Rexed’s laminae I−X is shown. Scale bar, 200 μm. (**C**) Left, synaptic connections between interneurons and motoneurons in the sham, CLES onset, untrained, and 10- to 20-Hz CLES groups. Scale bar, 20 μm. Middle, a three-dimensional model of the connection between interneurons and motoneurons. Scale bar, 20 μm. Right, higher-magnification images of the boxed area. Yellow regions (overlap of GFP and mCherry staining) indicate areas of connection between interneurons and motoneurons. Scale bar, 10 μm. (**D**) The contact area of interneurons per 1000 μm^2^ of motoneuron surface area in the sham, CLES onset, untrained, and 10- to 20-Hz CLES groups (n = 6 mice). (**E**) Bar plot showing the contact number of interneurons per 1000 μm^2^ motoneuron surface area in the sham, CLES onset, untrained, and 10- to 20-Hz CLES groups (n = 6 mice). Data are shown as the mean ± SEM, ns: no statistical difference, \**p* < 0.05, \*\**p* < 0.01, \*\*\**p* < 0.001, one-way ANOVA followed by Bonferroni’s post-test.

### Reconstruction of glutamatergic synaptic connections between neurons in SCI mice by 10- to 20-Hz CLES

We detected changes in the SCEP waveforms 1 week after SCI and found that there was no significant change in ER amplitude compared with the sham group, whereas the MR and LR amplitudes significantly decreased or disappeared (**Figure 9A, E, I**). These show that the sensorimotor circuit of the spinal cord was destroyed after SCI, which led to the blockade of polysynaptic electrical conduction.

**Figure 9.**
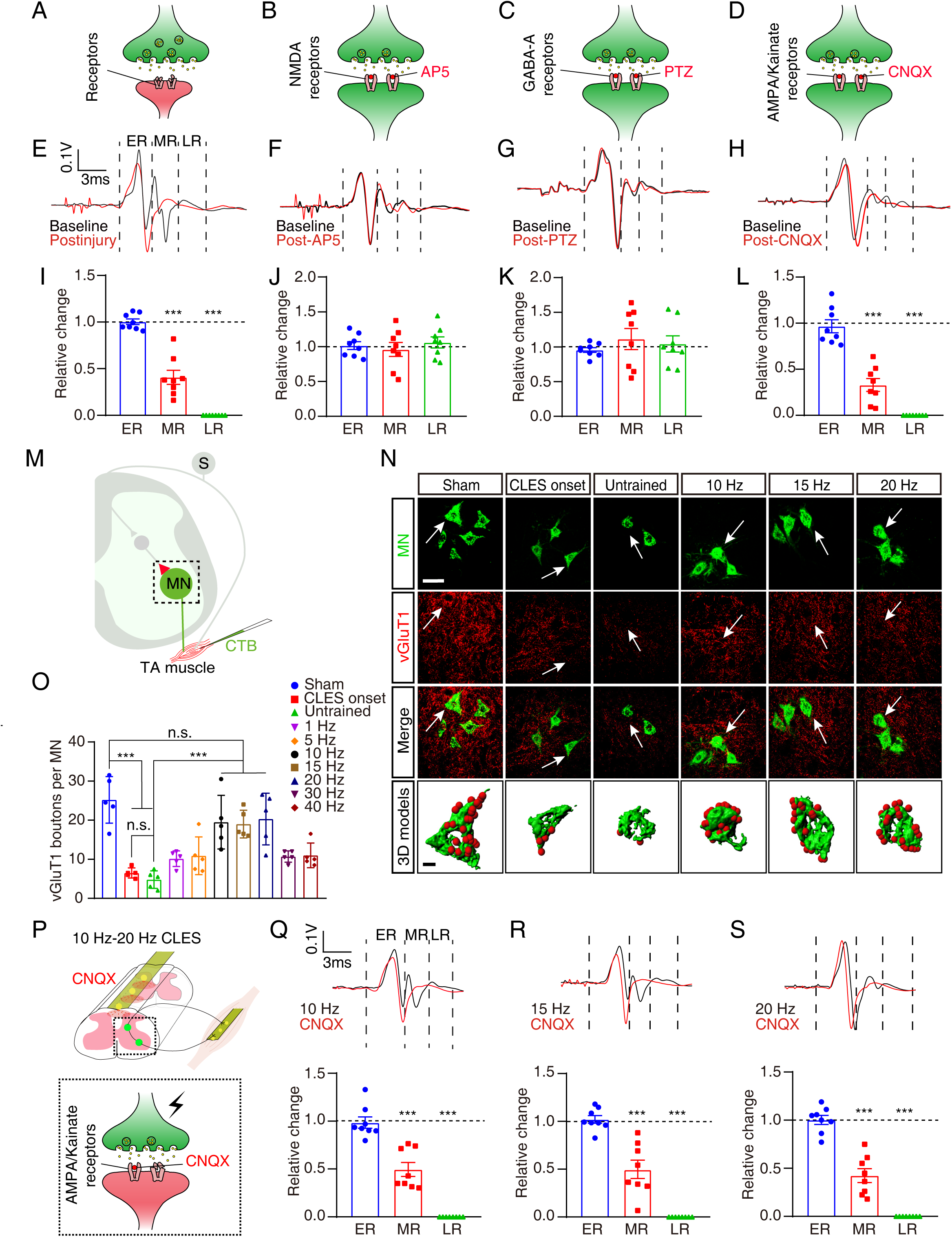
Remodeling of glutamate synaptic connections between spinal neurons in SCI mice after CLES. (**A**-**D**) Diagram of synaptic connections between spinal neurons after SCI (**A**) and after administration of AP5 (**B**), PTZ (**C**), or CNQX (**D**) in intact mice. (**E**) SCEPs were recorded before and 1 week after SCI. (**F**-**H)** SCEPs recorded before and 30 minutes after intrathecal injection of AP5 (**F**), PTZ (**G**), and CNQX (**H**) in intact mice. Each waveform was the average of responses to 50 stimuli. (**I**) The relative changes in ER, MR, and LR amplitude after SCI compared with the baseline of intact mice (n = 8 mice). (**J**-**L**), Compared with the baseline of intact mice, the relative change of ER, MR, and LR amplitude after intrathecal injection of AP5 (**J**), PTZ (**K**), and CNQX (**L**) (n = 8 mice). (**M**) Diagram illustrating how motoneurons were traced. (**N**) CTB was injected into the TA muscle to retrogradely trace motoneurons (green). vGluT1 (red) labeled axonal terminals of glutamatergic neurons in the sham, CLES onset, untrained, and 10- to 20-Hz CLES groups. Scale bar, 50 μm. The motoneuron and vGluT1 terminal indicated by the white arrow in each image was subjected to three-dimensional modeling. Scale bar, 10 μm. (**O**) The number of vGluT1 boutons in contact with each motoneuron (n = 5 mice). (**P**) Detecting SCEPs of SCI mice in the 10- to 20-Hz CLES groups after CNQX administration. The dashed box shows the synaptic connection between neurons after administration of CNQX. **(Q**-**S**) SCEPs were recorded (upper traces) in the 10-Hz (**Q**), 15-Hz (**R**), and 20-Hz (**S**) CLES groups before and after administration of CNQX. Bar plots (lower graphs) show relative changes in ER, MR, and LR amplitudes before and after CNQX administration (n = 8 mice). Data are shown as the mean ± SEM, ns: no statistical difference, \*\*\**p* < 0.001, Student’s *t*-test (**I**-**L** and **Q**-**S**), one-way ANOVA followed by Bonferroni’s post-test (**O**).

To clarify the receptors involved in the sensorimotor circuits blocked by SCI, we performed a series of pharmacological and electrophysiological tests. First, SCEPs were detected in intact mice; then, in the same mice, we blocked glutamate binding to N-methyl-D-aspartic acid (NMDA) receptors through local intrathecal delivery of AP5 at L5/L6 spinal segments (**Figure 9B**). Similarly, we administered intrathecal PTZ to other mice to prevent γ-aminobutyric acid (GABA) from binding to GABA-A receptors (**Figure 9C**). There was no significant change in ER, MR, or LR amplitudes after the use of AP5 and PTZ (**Figure 9F, G, J, K**). In contrast, we prevented glutamate from binding to AMPA/kainate receptors through local intrathecal delivery of CNQX at L5/L6 spinal segments, and the results showed that the amplitude of MR and LR of SCEPs was significantly decreased or disappeared (**Figure 9D, H, L**). Taken together, these results suggest that the glutamatergic synaptic connections between neurons in sensorimotor neural circuit, specifically those involving AMPA/kainate receptors were disrupted after SCI.

We then injected CTB into the TA muscle of mice after SCI to retrogradely trace the corresponding motoneurons in the spinal cord (**Figure 9M**). Vesicular glutamate transporter 1 (vGluT1) staining of motoneurons was performed in the same mice to label the glutamatergic axon terminals of premotor neurons (**Figure 9N**). Three-dimensional reconstruction of staining images was performed to quantify the vGluT1- positive boutons on motoneurons in each group (**Figure 9N and Figure 9-figure supplement 1**). Compared to sham group, the average number of vGluT1-positive boutons on each motoneuron was significantly reduced in the untrained and CLES onset group mice (**Figure 9O**). This indicated the disruption of glutamatergic synaptic connections between premotor neurons and motoneurons one week after SCI. The average number of vGluT1-positive boutons on each motoneuron was restored significantly in the 10- to 20-Hz CLES groups compared with the untrained group (**Figure 9O**). And, compared with the sham group, the average number of vGluT1- positive boutons on each motoneuron in the 10- to 20- Hz CLES groups almost recovered to the preinjury level (**Figure 9O**). These results indicate the reformation of glutamatergic synapses between premotor neurons and motoneurons after SCI.

The MRs and LRs of SCEPs in SCI mice were restored by 10- to 20-Hz CLES (**Figure 2A**). To verify recovery of the LR by 10- to 20-Hz CLES after SCI, glutamatergic synapses between premotor neurons and motoneurons were reconstructed. SCEPs were recorded in mice in the 10- to 20-Hz CLES groups after intrathecal injection of CNQX (**Figure 9P-S**). The ER amplitude did not change significantly after the application of CNQX, but the amplitude of the MR and LR was significantly decreased or disappeared (**Figure 9Q-S**). Together, our evidence demonstrates that 10- to 20-Hz CLES consistently promoted synapse reformation between glutamatergic neurons and motoneurons. This evidence also indicates that the spinal sensorimotor circuit was rebuilt morphologically with 10- to 20-Hz CLES after SCI.

### Activation of motoneurons by premotor neuron Input in SCI mice after 10- to 20-Hz CLES

To identify whether physiologically functional synaptic connections between premotor neurons and motoneurons could be reestablished by 10- to 20-Hz CLES in the sensorimotor circuit, Ca^2+^ signals from motoneurons of awake mice were recorded by using fiber photometry (**Figure 10A**). Retro-AAV-GCaMP6m was injected into the TA muscle, and the calcium indicator GCaMP6 was expressed in the corresponding motoneurons of the TA muscle. We implanted a small optical fiber (250 μm in diameter) with its tip in the motoneuron pool to record changes in GCaMP fluorescence. We first examined GCaMP signals when the sham group mice moved. Hindlimb movement reliably increased GCaMP fluorescence across the bouts of movement in these mice (**Figure 10B**). The enhancement of GCaMP fluorescence in motoneurons indicated that premotor neurons transmitted neural signals to motoneurons and activated them. Ca^2+^ signals were recorded in the motoneurons of mice under different experimental conditions (**Figure 10C-L**). SCI resulted in a significant reduction in Ca^2+^ signal intensity as compared with that of the sham group (**Figure 10M**). In contrast, the intensity of GCaMP signals in motoneurons were significantly stronger in 10- to 20-Hz CLES mice than in untrained mice (**Figure 10M**). In addition, the AUC of the motoneuron calcium signal change curve was measured. The average AUC decreased significantly after SCI (**Figure 10N**). The average AUC was significantly increased by 10- to 20-Hz CLES as compared with that of the untrained group (**Figure 10N**). And, the average AUC in the 10- to 20- Hz CLES groups almost recovered to a preinjury state (**Figure 10N**). In summary, these findings indicate that the premotor inputs of motoneurons in the sensorimotor neural circuits were destroyed by SCI, and the functional synaptic connections between premotor neurons and motoneurons were reconstructed by 10- to 20-Hz CLES.

**Figure 10.**
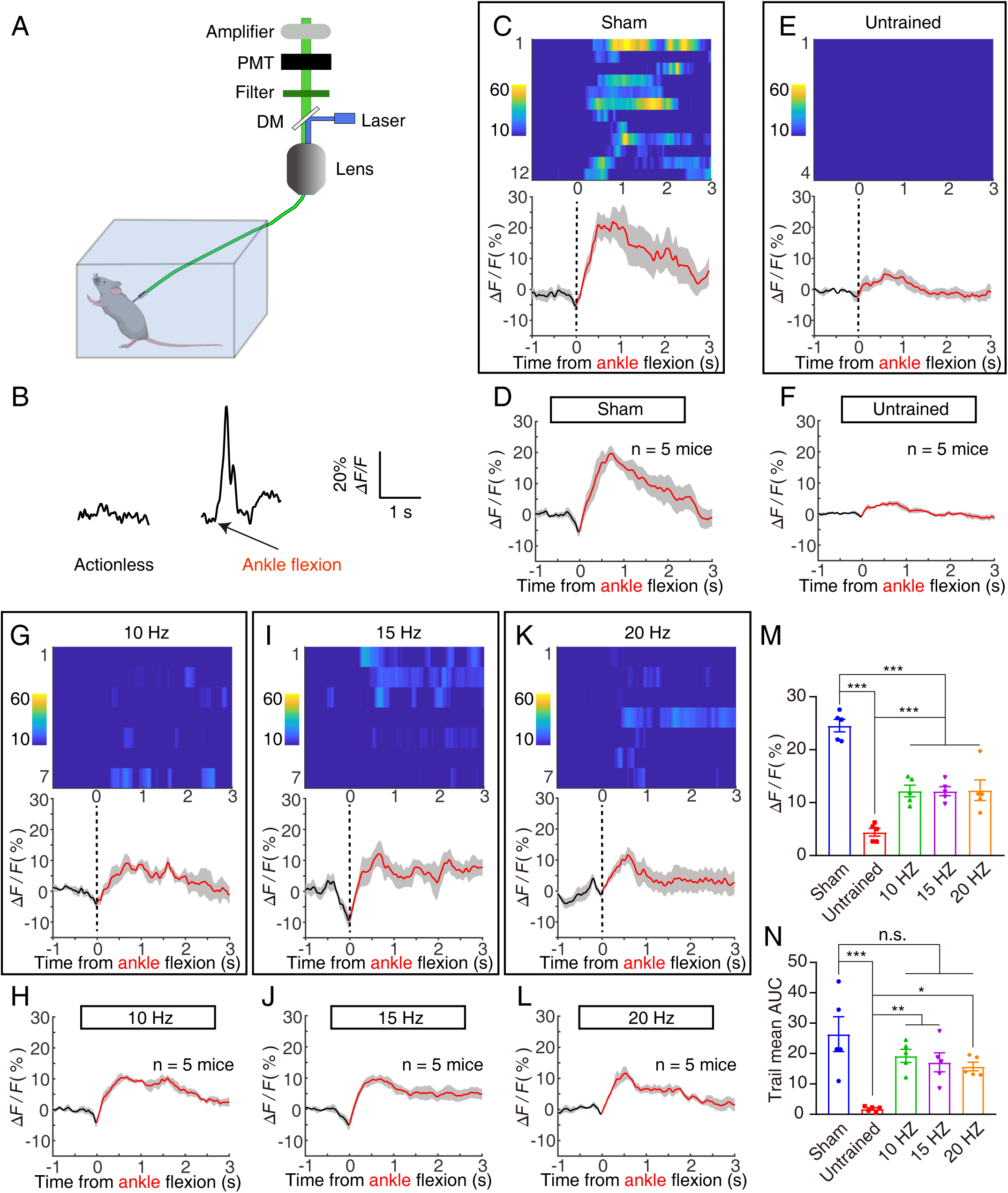
Recorded Ca^2+^ signals of lumbar motoneurons in SCI mice after CLES. (**A**) Schematic of the fiber photometry setup. Ca^2+^ transients were recorded from motoneurons of mice that had free movement. DM, dichroic mirror; PMT, photomultiplier tube. (**B**) Raw traces of GCaMP6 fluorescence changes that were related to flexion of the ankle joint of the hind limbs. Δ*F/F* represents the change in fluorescence from the mean level before the task. (**C**, **E**, **G**, **I**, and **K**) Ca^2+^ signals associated with ankle flexion in a single mouse from the sham (**C**), untrained (**E**), 10- Hz CLES (**G**), 15-Hz CLES (**I**), and 20-Hz CLES (**K**) groups. Upper, heatmap illustration of Ca^2+^ signals aligned with the initiation of ankle flexion bouts. Each row plots one bout; the number of bouts illustrated varies from 4 to 12. Color scale at the left indicates Δ*F/F*. Lower, plot of the average Ca^2+^ transients. Thick lines indicate the mean, and shaded areas indicate the SEM. The red line indicates the time of ankle flexion. (**D**, **F**, **H**, **J**, and **L)** Mean Ca^2+^ transients associated with ankle flexion for the entire test group (n = 5 mice) for each condition: sham (**D**), untrained (**F**), 10-Hz CLES (**H**), 15-Hz CLES (**J**), and 20-Hz CLES (**L**). (**M**) The average peak fluorescence of ankle flexion in the test chamber for 100 s (n = 5 mice). (**N**) The AUC of the average fluorescence peak of ankle flexion in the test chamber within 100 s (n = 5 mice). Data are shown as the mean ± SEM, ns: no statistical difference, \**p* < 0.05, \*\**p* < 0.01, \*\*\**p* < 0.001, one-way ANOVA followed by Bonferroni’s post-test.

### Characterization of molecular network changes in spinal motoneurons activated by 10- to 20-Hz CLES

To gain mechanistic insights into the reassembly of the spinal sensorimotor circuits with 10- to 20-Hz CLES, we performed gene expression profiling analysis in motoneurons. At 1 week postinjury, TRDA was injected into the TA muscle of mice to retrogradely trace the corresponding motoneurons in the spinal cord, and SCI mice were sacrificed after 3 weeks of 10- to 20-Hz CLES training. TRDA-labeled spinal motoneurons were isolated by the glass microtube aspiration method ^25^ under a fluorescence microscope (**Figure 11A**). mRNA from the purified spinal motoneurons was then used for library preparation and sequencing (**Figure 11A**). Transcriptome analysis identified 1,345 genes for which transcription had been altered (fold change ≥ 2.0; *p* < 0.05) in motoneurons from the 10- to 20-Hz CLES mice as compared with motoneurons from the untrained mice (**Figure 11-figure supplement 1A**). Among these genes, 559 were downregulated and 786 were upregulated (**Figure 11-figure supplement 1A**). Gene ontology (GO) analysis showed that one group of GO terms accounted for a large fraction of the upregulated transcripts (**Figure 11-figure supplement 1B**). This group includes axon growth−related cellular functions, such as cell adhesion, regulation of cell growth, and cell projection assembly (**Figure 11-figure supplement 1B**), suggesting that axon growth programs were reactivated with 10- to 20-Hz CLES. Then, we used qPCR to detect the mRNA expression levels of some genes in the lumbar spinal cord, and the results showed that the detected gene expression trends were consistent with the results of single-cell transcriptome sequencing (**Figure 11B, H**). These results support the idea that the axon growth program used during development was activated by 10- to 20-Hz CLES to promote remodeling of the spinal sensorimotor circuit after SCI.

**Figure 11.**
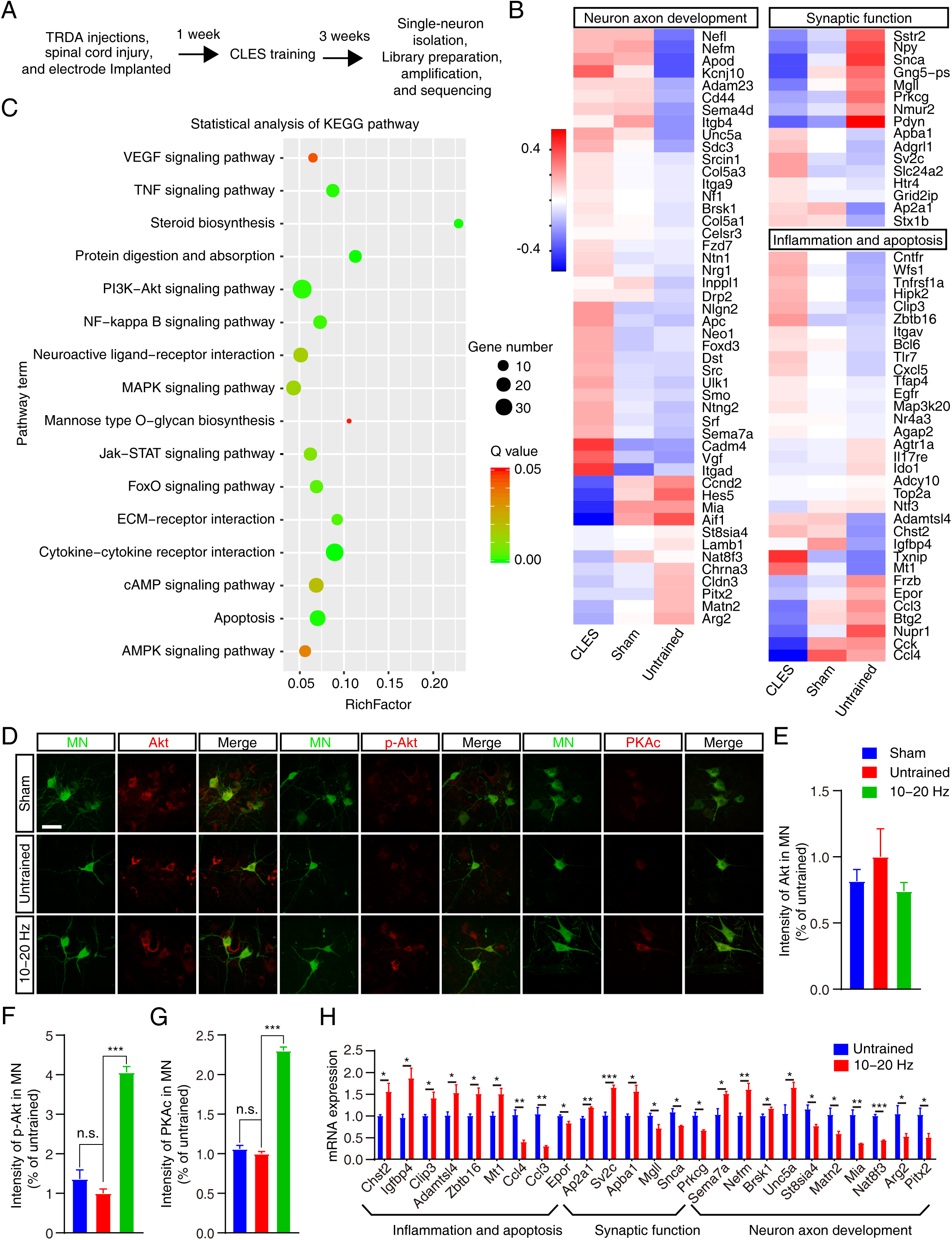
Changes in the transcripts of motoneurons in the spinal sensorimotor circuits of SCI mice after CLES. (**A**) Overview of RNA sequencing and analysis. (**B**) Heatmap showing the expression of a selection of differentially expressed genes among the sham group, untrained group, and 10-20 Hz CLES groups. (**C**) Statistical analyses of signaling pathways from the Kyoto Encyclopedia of Genes and Genomes, featuring the functional distribution of related genes and the related signaling pathways that reached significance in the 10-20 Hz CLES groups (*q* < 0.05). Enrichment analysis (richness factor) indicates the relative numbers of differentially expressed genes that were identified in our study relative to the total number of genes. Dot size is correlated with the level of expression of genes involved in specific signaling pathways. Dot color indicates the *q*-value, which ranged from 0 (green) to 0.05 (red). (**D**) Coimmunostaining of Akt, p-Akt, PKAc, and MN on coronal sections in the sham, untrained, and 10-20 Hz CLES groups (n = 5 mice). (**E**-**G**) Quantification of the expression levels of Akt (**E**), p-Akt (**F**), and PKAc (**G**) proteins in motoneurons. Scale bar, 50 μm. (**H**) The mRNA expression levels of some genes in **B** were verified by qPCR in the lumbar spinal cord (n = 4 mice). Data are shown as the mean ± SEM, ns: no statistical difference, \**p* < 0.05, \*\**p* < 0.01, \*\*\**p* < 0.001, one-way ANOVA followed by Bonferroni’s post-test (**E-G**), Student’s *t*-test (**H**).

To provide further insight into the mechanisms involved, we also used bioinformatics to identify the potential involvement of genes related to signaling pathways (**Figure 11C**). We analyzed functional pathways to determine how the differentially expressed genes could be coordinated to achieve protein-network transduction to mediate sensorimotor circuit remodeling. The results suggest that sixteen signaling pathways were involved, two of which have been reported to regulate axonal regeneration; this was accomplished by using the statistical analyses of the Kyoto Encyclopedia of Genes and Genomes (KEGG) pathways (Fig. 11c; PI3K-Akt signaling pathway, *q* < 0.01; cAMP signaling pathway, *q* < 0.05) ^26–29^. Immunohistochemistry revealed that Akt did not change in retro-AAV-GFP−labeled motoneurons of mice in the sham group, the untrained group, and 10- to 20-Hz CLES groups (**Figure 11D, E**). As compared with the untrained group, however, expression of phosphorylated (p-)Akt and PKAc were upregulated in the 10-20 Hz CLES groups, suggesting that PI3K-Akt and cAMP signaling pathways were activated (**Figure 11D- G**).

For a few of these isolated signaling pathways, there was, however, an uncertain association with axonal regeneration, including the pathway for VEGF signaling (*q* < 0.05), TNF signaling (*q* < 0.01), Steroid biosynthesis (*q* < 0.01), Protein digestion and absorption (*q* < 0.01), NF-κB signaling (*q* < 0.01), neuroactive ligand-receptor (*q* < 0.05), MAPK signaling (*q* < 0.05), Mannose type O−glycan biosynthesis (*q* < 0.05), Jak−STAT signaling (*q* < 0.01), FoxO signaling (*q* < 0.01), and ECM-receptor interaction (*q* < 0.01), as well as signaling pathways for cytokine-cytokine receptor interactions (*q* < 0.01), Apoptosis (*q* < 0.01), and AMPK signaling (*q* < 0.05) (**Figure 11C**). These data indicate that a number of well-studied and unknown signaling pathways were activated by 10- to 20-Hz CLES after SCI and were likely involved in the dynamic repair processes within the sensorimotor circuit.

## Discussion

We developed a closed-loop electrical stimulation (CLES) system targeting the mice lumbosacral spinal cord and TA muscle, which mimics sensory feedback and feedforward muscle contraction loops. We discovered the stimulus frequency of 10-20 Hz under closed-loop conditions was required for structural and functional reconstruction of spinal sensorimotor circuits. This discovery provides a mechanistic framework for the design of neuromodulation systems based on spinal cord stimulation to improve the recovery of sensorimotor functions following neurological disorders.

The sensorimotor reflex circuit we focus on is composed of TA muscle, sensory neurons, motor neurons and interneurons. Motor neurons dominate TA muscles and have a direct impact on motor function ^30, 31^. Interneurons are essential for producing reflex and rhythmic motor activity as well as many other activities ^24, 32, 33^. Sensory feedback systems based on sensory neurons constantly monitor the consequences of motor action ^24, 34^. Studies have shown that epidural electrical stimulation enhances the excitability of the spinal cord and sensory feedback information is essential for motor recovery below the injury level after SCI ^16, 18, 35^. Therefore, we established a closed-loop electrical neuromodulation system to enhance motor output and sensory feedback by combining central and peripheral synergistic electrical stimulation.

We found that the amplitudes of MR and LR were significantly increased by 10- to 20-Hz CLES compared to the untrained group (**Figure 2I, J**). The sensorimotor function of the hind limb by 10- to 20-Hz CLES was significantly improved in the SCI mouse compared to the untrained group (**Figure 3A, E and Figure 4J, K**). A quantitative analysis showed that the EMG amplitudes of bursts from TA muscle were significantly increased in the 10- to 20-Hz CLES groups, as compared with the untrained group (**Figure 3Q**). Taken together, the results demonstrated that polysynaptic electrical conduction in sensorimotor circuits and hindlimb sensorimotor function were restored by 10- to 20-Hz CLES. However, the mice of 10- to 20-Hz CLES groups did not return to the level of the sham group in polysynaptic electrical conduction and hindlimb sensorimotor function (**Figure 2-4**). For example, the amplitudes of MR and LR in the 10- to 20-Hz CLES groups only reached 48.4% and 59.1% of that of the sham group, respectively; the amplitudes of the EMG in the 10- to 20-Hz CLES groups reached only 19.1% of that of the sham group. Similarly, the contractile intensity of TA muscle in mice of the 10- to 20-Hz CLES groups only reached 61.9% of that of the sham group. We speculate that the recovery effect of CLES was greatly undermined because of the short training time and a single muscle stimulation.

In spinal sensorimotor reflex circuits, spinal interneurons play an important role in sensorimotor reflex circuits, which integrate information feedback from sensory neurons and then transmit motor instructions to activate or inhibit motoneurons to control muscle contraction ^36, 37^. We found that innervation of motor neurons by sensory neurons and interneurons was disrupted by SCI, whereas disrupted connections were rebuilt by 10- to 20-Hz CLES (**Figure 6F, G, I, J and Figure 8D, E**). But the maturity of these reconstructed synapses was not equivalent to those in the non-injured condition because the level of restored connectivity was lower than in the sham group (**Figure 6F, G, I, J and Figure 8D, E**). For example, the number and area of sensory-motor connections only reached 54.3% and 34.1% of that of the sham group, and the number and area of interneuron-motoneuron connections only reached 57.6% and 30.3% of that of the sham group, respectively. In addition, compared with the untrained group, the TA motor neurons of the mice in the 10- to 20-Hz CLES groups were significantly activated (**Figure 10M**); while the activation level of the TA motor neurons in the 10- to 20-Hz CLES groups only reached 49.8% of that of the sham group. These results suggested that 10-to 20-Hz CLES effectively remodeled synaptic connections between neurons in spinal neural circuits, but the number, area and function of synaptic connections between neurons did not return to preinjury levels. With an exception, we found that the average volume of motoneuron soma and dendrites and the number of glutamatergic synaptic connections between premotor neurons and motoneurons reached 93.9%, 85.9%, and 77.8% of that of the sham group, respectively (**Figure 7C, D and Figure 9O**).

The results of functional and structural analysis of neural circuits showed that CLES consisting of only a single muscle and spinal cord significantly remodeled neural circuits, but most of the detected indicators did not return to preinjury levels. Longer CLES treatments may further close the gap with the sham group, leading to more thorough sensorimotor reflex circuit reconstruction and better functional recovery. However, in this study, due to technical limitations such as electrode material and size, we were able to perform safe electrode transplantation in the mice for only 4 weeks, thus limiting longer periods of 10- to 20-Hz CLES application. In the future, we will consider further improving the material and production process of the electrodes so that they can be transplanted into the mice for longer periods of time.

Theoretically, each neuron has its own inherent neural signal firing frequency ^38^. Real-time and multimodal variable electrical stimulation can effectively improve the plasticity of the spinal neural circuits after SCI ^39, 40^, but it is not clear how these effective stimuli can be decoded. In this study, we used a spinal-muscle CLES system to obtain neural signal coding rules (with respect to certain parameters) that promote sensorimotor reflex circuit reconstruction. Only 10- to 20-Hz CLES was able to promote the correct reconstruction of sensorimotor reflex circuits in the spinal cord, and these restored sensorimotor functions persisted for 2 weeks after electrical stimulation ended (data not shown). The question arises as to whether this electrical stimulation frequency will also be applicable to the reconstruction of other types of spinal circuits. It is likely that neuromodulation therapies of SCI target distinct neural structures but share common principles. Such future discoveries will provide a theoretical framework for the broad advancement of neuromodulation therapies to improve function after SCI.

## Materials and Methods

### Key Resources Table

| Reagent type (species) or resource | Designation | Source or reference | Identifiers | Additional information |
| --- | --- | --- | --- | --- |
| strain, strain background (Mus Musculus) | ChAT-iCre | The Jackson laboratory | JAX#006410 |  |
| strain, strain background (Mus Musculus) | Lbx1-iCre | GemPharmatech | This paper | Genetic mice were used to label interneurons |
| strain, strain background (Mus Musculus) | C57BL/6J | SLAC Laboratory Animal Company |  |  |
| antibody | Anti-GFP (Goat polyclonal) | Abcam | Ab6662 | 1:200 |
| antibody | Anti-CTB (Rabbit polyclonal) | Invitrogen | PA125635 | 1:200 |
| antibody | Anti-mCherry (Rabbit polyclonal) | Abcam | Ab183628 | 1:200 |
| antibody | Anti-vGluT1 (Mouse monoclonal) | Merk Millipore | MAB5502 | 1:200 |
| antibody | Anti-NF (Mouse monoclonal) | Abcam | Ab82259 | 1:500 |
| antibody | Anti-α-BTX-555 (Mouse monoclonal) | Thermo Fisher Scientific | B35451 | 1:1000 |
| antibody | Anti-syn (Mouse monoclonal) | Abcam | Ab8049 | 1:500 |
| antibody | Anti-Akt (Rabbit monoclonal) | Cell Signaling technology | 4691s | 1:200 |
| antibody | Anti-p-Akt (Rabbit monoclonal) | Cell Signaling technology | 4060 | 1:200 |
| antibody | Anti-PKAc (Rabbit polyclonal) | Cell Signaling technology | 4782 | 1:200 |
| antibody | 488-conjugated, anti-Goat (Donkey polyclonal) | Abcam | ab150129 | 1:800 |
| antibody | 488-conjugated, anti-Rabbit (Goat polyclonal) | Abcam | Ab150077 | 1:800 |
| antibody | 647- | Abcam | ab150075 | 1:800 |
|  | conjugated,<br>anti-Rabbit<br>(Donkey<br>polyclonal) |  |  |  |
| antibody | 488-conjugated,<br>anti-Mouse<br>(Goat<br>polyclonal) | Abcam | ab150117 | 1:800 |
| antibody | 555-conjugated,<br>anti-Mouse<br>(Goat<br>polyclonal) | Abcam | ab150118 | 1:800 |
| antibody | 555-conjugated,<br>anti-Rabbit<br>(Goat<br>polyclonal) | Abcam | ab150078 | 1:800 |
| recombinant<br>DNA reagent | RV-N2C (G)<br>-ΔG-EGFP | BrainVTA | R03001 | |
| recombinant<br>DNA reagent | AAV2/9-EF1α-DIO-oRVG-WPRE-hGH-pA | BrainVTA | PT-0023 | |
| recombinant<br>DNA reagent | AAV2/2Retro-EF1α-<br>GCaMp6m-WPRE-hGH-pA | BrainVTA | PT-1700 | |
| recombinant<br>DNA reagent | AAV2/2Retro-<br>nEf1α-<br>mCherry-WPRE-pA | BrainVTA | PT-0099 | |
| recombinant<br>DNA reagent | AAV2/2Retro-<br>hSyn-EGFP-<br>WPRE-pA | Taitool<br>Bioscience | S0237-2R |  |
| recombinant<br>DNA reagent | AAV2/9-hSyn-DIO-mCherry | Brain Case | BC-0025 |  |
| recombinant<br>DNA reagent | Cholera toxin<br>B subunit | Thermo Fisher<br>Scientific | C34775 |  |
| chemical<br>compound,<br>drug | Dextran,<br>Texas Red | Thermo Fisher<br>Scientific | D3328 |  |
| chemical compound, drug | D(-)-2-amino-5-phosphonopen tanoic acid | Sigma | A8054 |  |
| chemical compound, drug | pentylenetetra zole | Sigma | P6500 |  |
| chemical compound, drug | 6-cyano-7-nitroquinoxali ne-2,3-dione | MedChemExpres s | HY-15066 |  |
| software, algorithm | GraphPad Prism 8.0 | GraphPad Software |  |  |
| software, algorithm | Origin 2018 | OriginLab |  |  |
| software, algorithm | Imaris 0.0.1 | Oxford Instruments |  |  |
| software, algorithm | ZEN 2012 | ZEISS |  |  |
| software, algorithm | MATLAB | Mathworks |  |  |

### Animals

All the procedures were in accordance with the Institute of Neuroscience (Soochow University) guidelines for the use of experimental animals and were approved by the Institutional Animal Care and Use Committee at Soochow University. *ChAT^cre^* (JAX stock #006410) mouse strains were maintained on a mixed genetic background (129/C57Bl6), kindly provided by Dr Zilong Qiu (Institute of Neuroscience, Chinese Academy of Sciences, Shanghai, China). *Lbx1^cre^* transgenic mice were constructed by GemPharmatech Company. Wild-type C57BL/6J mice were purchased from the Shanghai SLAC Laboratory Animal Company (Shanghai, China). All mice including wild-type C57/BL6J, *ChAT^cre^*, and *Lbx1^cre^* mice were housed in the specific pathogen free animal facility at Soochow University. Mice were maintained on a 12/12 light/dark cycle and housed in groups of five for 8 weeks. After surgery, mice were housed individually for 1 week before further experiments. Mice determined to unhealthy or died were excluded (< 10%). Spinal T9 complete transection was judged to be successful when BMS score < 1 in mice on SCI 7d. All mice received analgesic (carprofen, 5 mg/kg) before wound closure and every 12 h for at least 48 h after injury. The health of animals was monitored daily for at least ten days after surgery, after which weekly health checks were performed.

### Electrode Implantation

All experimental mice were anesthetized with pentobarbital (40 mg/kg, i.p.). First, a laminectomy was performed at vertebral level T9 to create an entry point for the implant. An insertable epidural needle electrode plate for mice was slowly inserted along the gap between the exposed spinal cord and vertebrae so that metal contacts covered the L2−L4 segment of the spinal cord, and electrophysiological testing was performed intraoperatively to fine-tune the positioning of electrodes during this process. Ears on the electrode were sutured to the muscle to affix the electrode, and then the wound was sutured. Second, to record SCEP activity, an implantable electrode plate was implanted into the right hindlimb TA muscle after exposing that muscle. Then, the skin was sutured, and the electrode was fixed. Experiments were performed by an independent researcher who was blinded to the group allocation.

### Spinal Cord Injury

A spinal cord transection was performed during the implantation of the electrodes. After the spinal cord electrodes were implanted, and the spinal cord was completely transected at T9 using microscissors as described. Complete transection lesions were verified postmortem by confirming the absence of neural tissues throughout the dorsoventral extent of the spinal cord ^41, 42^. Sham operations were performed as follows: the dorsal vertebral lamina at spinal T9 segment was removed, the electrode was implanted to the spinal cord, but the spinal cord itself was left intact. After these operations, the muscle layers, fascia, and the skin were. Urine was expressed by manual abdominal pressure twice daily until mice regained reflex bladder function. Surgeries were performed by an independent surgeon who was blinded to the group allocation.

### Electric Stimulus Parameter

We found that motoneurons innervating the TA muscle were mainly distributed in the L2-L4 segment of the spinal cord, so that a spinal cord electrode was implanted to cover there and a muscle electrode was implanted in the TA muscle. A single pulse (0.2 ms) of spinal cord stimulation elicited spinal cord evoked potentials (SCEPs) in TA muscles; LR amplitudes of SCEPs reached a peak when the spinal cord stimulation intensity was 0.5-0.7 V, indicating that feedforward transmission mediated by spinal sensorimotor circuits was fully activated. Therefore, the spinal cord stimulation intensity was adjusted between 0.5 V and 0.7 V in the CLES and OLES-SCS. Spinal epidural recordings confirmed an effective response (unimodal peak) with minimal stimulation of 1 V administered to the TA muscle. Thus, TA muscles received a threshold electrical stimulation of 1V in the CLES and OLES-MS systems, transmitting sensory feedback signals to the spinal cord.

It has been reported that 1-10 V spinal cord electrical stimulation promotes hind limb movement in SCI rats ^43–45^. We found that the physiological response of mice was unstable when the spinal cord electrical stimulation intensity was higher than 2 V, so the intensity of the spinal cord electrical stimulation was 2 V in the OLES-SCShv group.

Experiments were performed by an independent researcher who was blinded to the group allocation.

#### Experimental Groups

In all experiments and groups, the spinal cord electrode and the muscle electrode were implanted as mentioned in “Electrode Implantation”. In the Sham group, the spinal cord was intact and electrical stimulation training was not implemented. In the Untrained group, the spinal cord was completely transected and electrical stimulation training was not implemented.

### CLES

Awake mice with complete spinal cord transection were fixed in a simple fixation device for electrical stimulation, and a programmable small-animal motor function rehabilitation training device (Suzhou Institute of Biomedical Engineering and Technology, Chinese Academy of Sciences, Suzhou, Jiangsu, China) was connected to the spinal cord stimulation electrode and the muscle stimulation electrode for stimulation training. Electrical stimulation parameters are described above. The spinal cord and muscles were given positive and negative rectangular pulses (duration, 0.2 ms) at 50 ms intervals with a 10 minutes rest period after every 15 minutes stimulation training. Different experimental groups were trained with different stimulus frequencies for a total of 3 weeks, with training taking place 6 days per week. Mice in the CLES group were trained daily for 90 minutes, starting from 7 days after SCI.

### OLES-SCS

The stimulation conditions of the group are the same as the CLES group except that muscle stimulation was not performed.

### OLES-MS

The stimulation conditions of the group are the same as the CLES group except that spinal cord was not performed.

### OLES-SCShv

The stimulation conditions of the group are the same as the *OLES-SCS* group except that the voltage intensity of spinal cord stimulation was 2 V.

Experiments were performed by an independent researcher who was blinded to the group allocation. Mice were randomly assigned to each group to receive training.

### SCEPs recording

In anesthetized mice, a 16-channel physiological signal recording and analysis system (BIOPAC, MP150, USA) was used to record SCEPs. The spinal cord stimulation electrode was connected to the STM 100C stimulation module, and the muscle electrode was connected to the EMG 100C recording module. AcqKnowledge software (BIOPAC, USA) was used to set the parameters of the STM 100C stimulation module, including a stimulation pulse frequency of 1 Hz and a pulse width of 0.2 ms. When the EMG 100C recording module was used to collect signals, the following filter settings were used: Band Stop channel 1 parameter, 50 Hz; Band Stop channel 2 parameter, 100 Hz; and Low Pass channel parameter, 2000 Hz.

Fifty SCEPs were recorded in each mouse, and data were exported in TXT format. The MATLAB software (MathWorks, Inc., USA) program for analyzing SCEPs, which was developed in collaboration with Dr. Shouyan Wang (Institute of Brain-like Intelligent Science and Technology, Fudan University, Shanghai, China), was used to divide and align the data. A series of continuous SCEPs were divided into single waveforms of the same duration, and then fifty single waves were aligned and superimposed. SCEPs baselines were calibrated, and waveforms were averaged to calculate the peak value and the AUC of the waveform.

Experiments were performed by an independent researcher who was blinded to the group allocation.

### Electrophysiological and Pharmacological Experiments

Pentobarbital (40 mg/kg, i.p.) was injected to anesthetize the mice implanted with electrodes. The following experiments were then conducted: (1) EES was applied to the spinal cord using a 1-Hz frequency and 0.2 ms pulse width so that SCEPs were recorded in the TA muscle, and (2) pharmacological modulation of EES-evoked motor responses was tested using the sodium channel blocker tetrodotoxin (TTX; 1 μM, MedChemExpress, USA), which blocks synaptic transmission; the 2-adrenergic receptor agonist tizanidine (1 mM, MedChemExpress), which decreases the excitability of spinal interneurons supplied by Group II fibers; pentylenetetrazole (PTZ; 5 mM, Sigma, USA), an inhibitor of γ-aminobutyric acid (GABA) and the GABA-A receptor; D(−)-2-amino-5-phosphonopentanoic acid (AP5; 0.5 mM, Sigma), an NMDA receptor antagonist; and 6-cyano-7-nitroquinoxaline-2,3-dione (CNQX; 50 μM, MedChemExpress), a potent and competitive AMPA/kainate receptor antagonist. All drugs (10 μl) were delivered intrathecally to the L5−L6 spinal segment. Experiments were performed by an independent researcher who was blinded to the group allocation.

### Spinal Injections

For spinal cord−targeted viral deliveries, we performed stereotaxic injections using high-precision instruments (Stoelting) while the mice were under pentobarbital anesthesia. The vertebrae of the L2−L4 segment of the spinal cord were exposed. A small hole was drilled in the middle of each of the three vertebrae in a rostral-to-caudal orientation, and a pulled calibrated glass pipette (Hamilton, USA) was used for local infusion of 500 nl of a virus solution (see below) by using a single-channel microinjection-withdrawal pump (KDS). The glass pipette was retracted after a 5 minutes period. The coordinates used to locate the spinal cord were 35.5 degrees from the midline of the spinal cord and a depth of 1 mm. Experiments were performed by an independent researcher who was blinded to the group allocation.

### Muscle Injections

The TA muscle was selected as the target muscle for injection of virus or tracer. A pulled calibrated glass pipette and a microsyringe were connected by using hot-melt adhesive. Virus or tracer was mixed with an appropriate amount of fast green dye (Sangon Biotech, China) before injection to facilitate observation. The TA muscle was divided into three regions from top to bottom, and 2 μl of virus or tracer was slowly injected into each of these regions (i.e., three injection sites); the glass pipette was retracted from each injection site after a 1 min period. Muscle injection specificity was always verified postmortem by the presence of fluorescence exclusively located in the targeted muscle. Experiments were performed by an independent researcher who was blinded to the group allocation.

### Spinal Sensorimotor Circuit Tracing

We labeled multiple elements of the spinal sensorimotor circuits: the motoneurons, the premotor circuit, proprioceptive axons, dorsal root ganglion (DRG) fibers, and sensory neuron−motoneuron connections. For retrograde labeling of motoneurons, we injected Retro-AAV-EGFP (Taitool Bioscience, China) into TA muscles as described above and perfused the mice 2 weeks later. The same injection method was used for CTB (Thermo Fisher Scientific, USA), and RV-N2C (G)-ΔG-EGFP (BrainVTA, China) delivery.

For G-deleted monosynaptic rabies premotor circuit tracing, rAAV-Ef1α-DIO-RVG-WPRE-pA (AAV-DIO-RVG; BrainVTA) was injected first into the ventral horn of the spinal cord of *ChAT^cre^*mice. Four weeks later, RV-N2C (G)-ΔG-EGFP was injected into the TA muscle.

CTB was injected into the TA muscle to anterogradely trace the corresponding proprioceptive axon in the spinal cord. Mice were sacrificed 4 weeks after injection.

To label DRG fibers, we pressure-injected 0.5 µl of pAAV-CAG-EGFP-3xFLAG-WPRE (AAV-GFP; BrainVTA) into the DRG at lumbar region L2−L4 using a finely pulled glass micropipette (at a coordinate of 0.3 mm below the DRG surface). Viral injections were performed 14 days prior to sacrifice. The micropipette was left in place for 3 minutes after the injection to avoid backflow. Finally, for all relevant procedures, the wounds were closed with layered sutures.

Approximately 2 µl of 10% Texas Red−dextran amine (TRDA; Invitrogen, USA) dissolved in ddH_2_O was slowly injected into TA muscles using glass micropipettes to mark specific motoneurons. At the same time, sensory nerve fibers were labeled as described above.

Experiments were performed by an independent researcher who was blinded to the group allocation.

### Tissue Clearing Technique

We first prepared the two CUBIC (clear, unobstructed brain / body imaging cocktails and computational analysis) solutions. Reagent 1 consisted of 12.5 g urea, 12.5 g *N*,*N*,*N*’,*N*’-tetramethylethylenediamine, and 17.5 ml water. This mixture was heated and stirred at 61 °C until completely dissolved; then, the solution was cooled to room temperature. Finally, 7.5 g Trion X-100 was added, and the solution was thoroughly stirred. Reagent 2 consisted of 25 g sucrose, 12.5 g urea, and 7.5 ml water. This mixture was heated and stirred at 61 °C until completely dissolved; then, the solution was cooled to room temperature. Finally, 5 g 2,2’,2’’-nitrilotriethanol and 400 μl of 10% Trion X-100 were added to the solution, and the solution was thoroughly stirred. These solutions were prepared fresh before each experiment.

The mice were perfused using standard methods, the spinal cord was removed and immersed in 4% paraformaldehyde (PFA; Sigma, USA) in refrigerator at 4 °C for 12 hours before proceeding to tissue clearing. To clear the tissue, the spinal dura mater of the L2−L4 segment was removed, and the tissue was transferred to a 50 ml centrifuge tube with 40 ml reagent 1 and placed in a 37 °C shaker for 4 days. The spinal cord was removed from reagent 1 and washed with phosphate-buffered saline (PBS; pH 7.4; Sinoreagent, China) with three changes over a 3 hours period, placed in 20% sucrose (Sinoreagent) in PBS for dehydration for 12 hours at 4°C, then, the tissue was transferred to a 50 ml centrifuge tube with 40 ml reagent 2 and placed in a black box at room temperature for 4 days.

Experiments were performed by an independent researcher who was blinded to the group allocation.

### Whole-Mount Immunohistochemical Staining

The spinal cord was placed in 20% sucrose (Sinoreagent) in PBS for dehydration for 12 hours at 4 °C (see above), and then embedded in O.C.T. (Sakura Finetek, Japan) in a −80 °C freezer for 12 hours. The next day, the tissue was washed with PBS (with three changes over a 3 hours period) and placed in goat anti-GFP (Ab6662; Abcam, USA) solution diluted with 2% BSA (Sangon Biotech) (1:50) for 3 days at room temperature. The tissue was then washed with 0.1% Trion X-100 in PBS (with three changes over a 3 hours period) and subsequently incubated with an appropriate secondary antibody (see below) and washed in the same way. Next, the tissue was transferred to 20% sucrose in PBS and was dehydrated overnight at room temperature. Finally, the tissue was placed in reagent 2. Experiments were performed by an independent researcher who was blinded to the group allocation.

### Immunohistochemistry

Pentobarbital (40 mg/kg, i.p.) was injected to anesthetize the mice before sacrificing them. Mice were perfused with 4% PFA in PBS. spinal cords were dissected out and washed with PBS. Then these tissues were dehydrated in 30% sucrose overnight to replace PFA with sucrose. After that, L2-L4 segement of spinal cord were embedded with O.C.T. for cryosectioning. After cryosectioning, coronal sections (35 μm thick) of the spinal cord were used for immunostaining. Primary antibodies diluted in PHT were incubated at 4 °C at the following concentrations: Go-GFP (Ab6662; Abcam), 1:200; Rb-CTB (PA125635; Invitrogen), 1:200; Rb-mCherry (ab183628; Abcam), 1:200; Mo-vGluT1 (MAB5502; Merck Millipore, USA), 1:200; Rb-Akt (4691s; Cell Signaling Technology, USA), 1:200; Rb-p-Akt (4060; Cell Signaling Technology), 1:200; Rb-PKAc (4782; Cell Signaling Technology), 1:50; Mo-NF (ab82259; Abcam), 1:500; α-BTX-555 (B35451; Thermo Fisher Scientific), 1:1000; Rb-syn (ab16659; Abcam), 1:500. The tissues were washed in PBS, incubated with the appropriate Alexa Fluor−conjugated secondary antibodies (1:800; Abcam) diluted in PHT at 4℃ for 12 hours, and washed with PBS again. Images were acquired with a Zeiss LSM700 confocal microscope (Carl Zeiss, Germany) and processed and exported with Zen software (Carl Zeiss). Experiments were performed by an independent researcher who was blinded to the group allocation.

### Motoneuron Reconstructions and Quantification

Spinal cord images were acquired using a LSM700 confocal microscope (Zeiss, 20× objective) in Z-stack mode, and at least 35 layers of pictures for each spinal cord section were taken. Motoneurons were reconstructed using IMARIS (Bitplane, Switzerland) software. Neuronal surfaces, including the unambiguously assigned dendrites, were reconstructed using the IMARIS surface tool before the overall surface area was calculated for each reconstructed neuron. The values in the text represent the mean volume and intensity of the reconstructed neurons and dendrites in each field of view. In each experimental group (n = 6 mice), the visual field with the largest number of neurons in the spinal cord slice was selected to be imaged. Quantifications were performed by an independent researcher who was blinded to the group allocation.

### Analysis of Interneurons, Dorsal Root Ganglion (DRG) Fibers, and Motoneuron Connections

Images were acquired with a Zeiss LSM700 confocal (64× objective) microscope in Z-stack mode. Complete motoneurons or dendrites were continuously scanned. The IMARIS contact area tool was used to assess the synaptic input of interneurons and DRG fibers to motoneurons. Quantifications were performed by an independent researcher who was blinded to the group allocation.

### Synaptic Analysis

High-resolution analysis of glutamatergic synaptic input to retrogradely marked motoneurons was acquired with a confocal (Zeiss, 64× objective) microscope in Z-stack mode and quantified using IMARIS software. We used the IMARIS point detection function to quantitatively distinguish vGluT1 labeling with a diameter of 2 μm as an estimate of glutamatergic neuron input. The ‘find spots close to surface’ option in the spot detection function was selected to quantitatively distinguish the effective connection between glutamatergic neurons and motoneurons. For proprioceptive axon input analyses to the spinal cord, we used the IMARIS spot detection function to quantitatively differentiate axon inputs > 1.5 μm in diameter as an estimation of proprioceptive input. Quantifications were performed by an independent researcher who was blinded to the group allocation.

### Analysis of Dendritic Complexity

A LSM700 confocal (Zeiss, 20× objective) microscope was used to acquire images of the spinal cord in the L2−L4 segment in Z-stack mode, and at least 200 layers of pictures were taken per mouse. The area with the largest number of motoneurons was selected, and the Filament Tracing module in IMARIS was used to reconstruct the dendritic morphology of motoneurons in that area. The number of neurons and the total length of dendrites were counted, and the average dendrite length of each motoneuron was calculated to measure the complexity of the neuron. Quantifications were performed by an independent researcher who was blinded to the group allocation.

### Myofilament and Neuromuscular Junction (NMJ) Analyses

The TA muscle was fixed with 4% PFA in a refrigerator at 4 ℃ and washed three times in 0.01 M PBS for 10 minutes each time. Subsequently, the TA muscle was dehydrated by sucrose, embedded in O.C.T. Hematoxylin and eosin were used to stain the frozen sections of tissue for histological analysis. Images were acquired with a Zeiss microscope (20× objective) and processed and exported with Zen software. Finally, the area of each myofilament was measured by ImageJ (NIH), and Graph Prism 8 (Graph Pad, La Jolla, CA) was used to analyze the differences among groups ^46^.

To stain the NMJs, the TA muscle was fixed with 4% PFA in a refrigerator at 4℃ for 12 hours. After being washed in PBS, the muscle was separated into bundles of 5−10 muscle fibers under a microscope and placed in PHT for 1 hour at room temperature. The antibodies Mo-NF (ab82259, Abcam; 1:500), Rb-syn (ab16659, Abcam; 1:500), and α-BTX (B35451, Thermo Fisher Scientific; 1:1000) were used to label the NMJs. Images were acquired using the LSM700 confocal microscope in Z-stack mode. The percentage of pre-synaptic (syn) area to post-synaptic (α-BTX) area was calculated in each neuromuscular junction ^47^.

Experiments were performed by an independent researcher who was blinded to the group allocation.

### Fiber Photometry

Retro-AAV-GCaMP6m virus (BrainVTA) was injected into TA muscle to express GCaMP in the corresponding motoneurons in the spinal cord. Four weeks later, vertebrae in the L3 segment of the spinal cord were exposed, and the right side of vertebrae of the spinal cord were opened with a skull drill. An optical fiber (250 μm O.D., 0.37 NA; Shanghai Fiblaser) was placed in a ceramic ferrule. The ceramic ferrule was attached to the stereotaxic instrument by a special holder, and then the optical fiber was implanted in the spinal cord (0.3 mm to the right of the midline of the spinal cord at an angle of 20 degrees; implantation depth, 0.7−0.8 mm). Biological tissue glue and dental cement were used to attach the ceramic ferrule to the vertebra. Mice were individually housed for at least 1 week to recover.

To record fluorescence signals, an optical fiber (250 μm O.D., 0.37 NA, 2 m long) guided the light between a commutator (Doric Lenses) and the implanted optical fiber. The laser power was adjusted such that the light emitted at the tip of the optical fiber was at a low level of 10−30 μW to minimize bleaching. Analog voltage signals were digitized at 200−500 Hz and recorded by CamFiberPhotometry software (Thinker Tech Nanjing Biotech Limited, China).

After 4 weeks of GCamP6m virus expression, calcium signals were recorded while the mice moved freely. The timing of behavioral variables was recorded with the same system. The behavioral processes and calcium signals were synchronized and analyzed by MATLAB software.

To calculate the GCamP6m signal, the relative fluorescence changes of ΔF/F were calculated to determine the Ca^2+^ signal as follows:

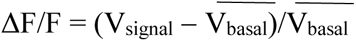

where 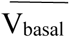 is the average value of V_basal_ in reference time and V_signal_ is the fluorescent signal collected during the experiment.

Experiments and quantifications were performed by an independent researcher who was blinded to the group allocation.

### Basso Mouse Scale (BMS) Scoring

Mice were tested for hindlimb functional deficits at 0, 1, 3, 7, 14, 21, and 28 days (n = 10 per group) after SCI. Hindlimb locomotor recovery was assessed in an open field using the BMS, which was previously described in detail ^48^. This scale ranges from 0, indicating complete paralysis, to 9, indicating normal hindlimb movement. Quantifications were performed by an independent researcher who was blinded to the group allocation.

### Electromyography (EMG) Recordings

Starting 4 weeks postinjury, training took place on a body weight−supporting (BWS, ∼60–80%, 2 m·minute^−1^) treadmill (SANS, China). EMG and behavioral recordings were started once the animals had achieved stable locomotor performance. A multichannel electrophysiological recording system (BIOPAC MP150) was used to record the EMG signal in TA muscle. Experiments were performed by an independent researcher who was blinded to the group allocation.

### Quantitative Evaluation of Motor and Sensory Function

After the 3 weeks CLES and OLES training protocol, the muscle electrode was removed from the mice, and the skin was sutured. A pressure-sensitive sensor was attached to the skin overlying the TA muscle, and the contractile function of the TA muscle was detected as described ^49^.

Mice were tested for hindlimb sensory recovery at 0, 7, 14, 21, and 28 days (n = 8 per group) after SCI by 10 up-down von Frey filament tests. Briefly, mice were placed under a small, opaque container on an elevated wire mesh. Animals were acclimated to the testing environment for 30 minutes prior to testing. Hind paws were assessed using the up-down method, and each hind paw received 10 consecutive trials to achieve a 50% withdrawal threshold as described ^50^.

Quantifications were performed by an independent researcher who was blinded to the group allocation.

### Single-Cell Transcriptome Analysis

Spinal cord motoneurons of mice in the sham group (n = 3 mice), untrained group (n = 3 mice), 10−20 Hz CLES group (10 Hz, n = 1 mice; 15 Hz, n = 1 mice; 20 Hz, n = 1 mice), were labeled with TRDA and sacrificed 4 weeks after SCI. The freshly isolated spinal cord was sliced on a Vibratome (Leica, Germany) into 200- to 300-μm-thick coronal sections. Fluorescence-labeled motoneurons were visualized under a fluorescence microscope (Zeiss) using a miniature operating system (Eppendorf, Germany), and the motoneurons were aspirated by glass electrodes and placed in preservation solution (Invitrogen, USA).

TRDA-labeled neurons were individually homogenized in 1 ml TRIzol (Invitrogen), and RNA was extracted with a RNeasy Mini Kit (Qiagen) or equivalent. Total RNA (1 μg) was used to synthesize double-stranded cDNA. First, reference genome sequences and gene model annotation files of related species were downloaded from genome websites such as UCSC, NCBI, and ENSEMBL. Second, Hisat2 software (v2.0.1, http://daehwankimlab.github.io/hisat2/) was used to index the reference genome sequence. Finally, clean data were aligned to the reference genome via Hisat2 software (v2.0.1). Then, transcripts in FASTA format were converted from known gff annotation files and indexed properly. Then, with the file that served as the reference genome file, HTSeq (v0.6.1, https://htseq.readthedocs.io/en/master/) was used to estimate gene and isoform expression levels from the paired-end clean data. Differential expression analysis was performed using the DESeq2 Bioconductor package, a model based on a negative binomial distribution. The estimates of dispersion and logarithmic fold changes incorporated data-driven prior distributions ^51^, and the adjusted *p*-value of the genes was set to <0.05 to detect differentially expressed genes. GOSeq (v1.34.1, http://bioconductor.org/packages/release/bioc/html/goseq.html) was used to identify gene ontology (GO) terms that were associated with the list of enriched genes with a significant adjusted *p*-value OR padj of <0.05. In addition, topGO was used to plot directed acyclic graph ^52^. Kyoto Encyclopedia of Genes and Genomes (KEGG) is a collection of databases associated with genomes, biological pathways, diseases, drugs, and chemical substances (http://en.wikipedia.org/wiki/KEGG). We used in-house scripts to isolate significantly differentially expressed genes that were represented in the KEGG database.

Experiments and quantifications were performed by an independent researcher who was blinded to the group allocation.

### RNA Isolation and Quantitative Real-Time Polymerase Chain Reaction

Total RNA was extracted using Tissue RNA Purification Kit Plus (YiShan Biotechnology, China) according to the protocol. Total RNA (1 μg) was used to synthesize cDNA by reverse transcription using HiScript III All-in-one RT SuperMix Perfect for qPCR Kit (Vazyme, China). Quantitative real-time polymerase chain reaction (qPCR) test was conducted by 2× SYBR Green qPCR Master Mix (Low ROX) (Bimake, USA) using real-time PCR Detection System (ABI 7500, Life technology, USA). The cycling conditions included a 10 minutes initial denaturation step at 95℃ followed by 40 cycles of 15 s at 95℃, 30 s at 60℃ and 30 s at 72℃. Target gene expression was normalized to the housekeeper gene GAPDH expression. Relative fold difference in expression was calculated using 2^−ΔΔCt^ method after normalization to GAPDH expression. The primers (Genewiz and Sangon Biotech, China) are shown in **Table 1**. Experiments and quantifications were performed by an independent researcher who was blinded to the group allocation.

**Table 1.**
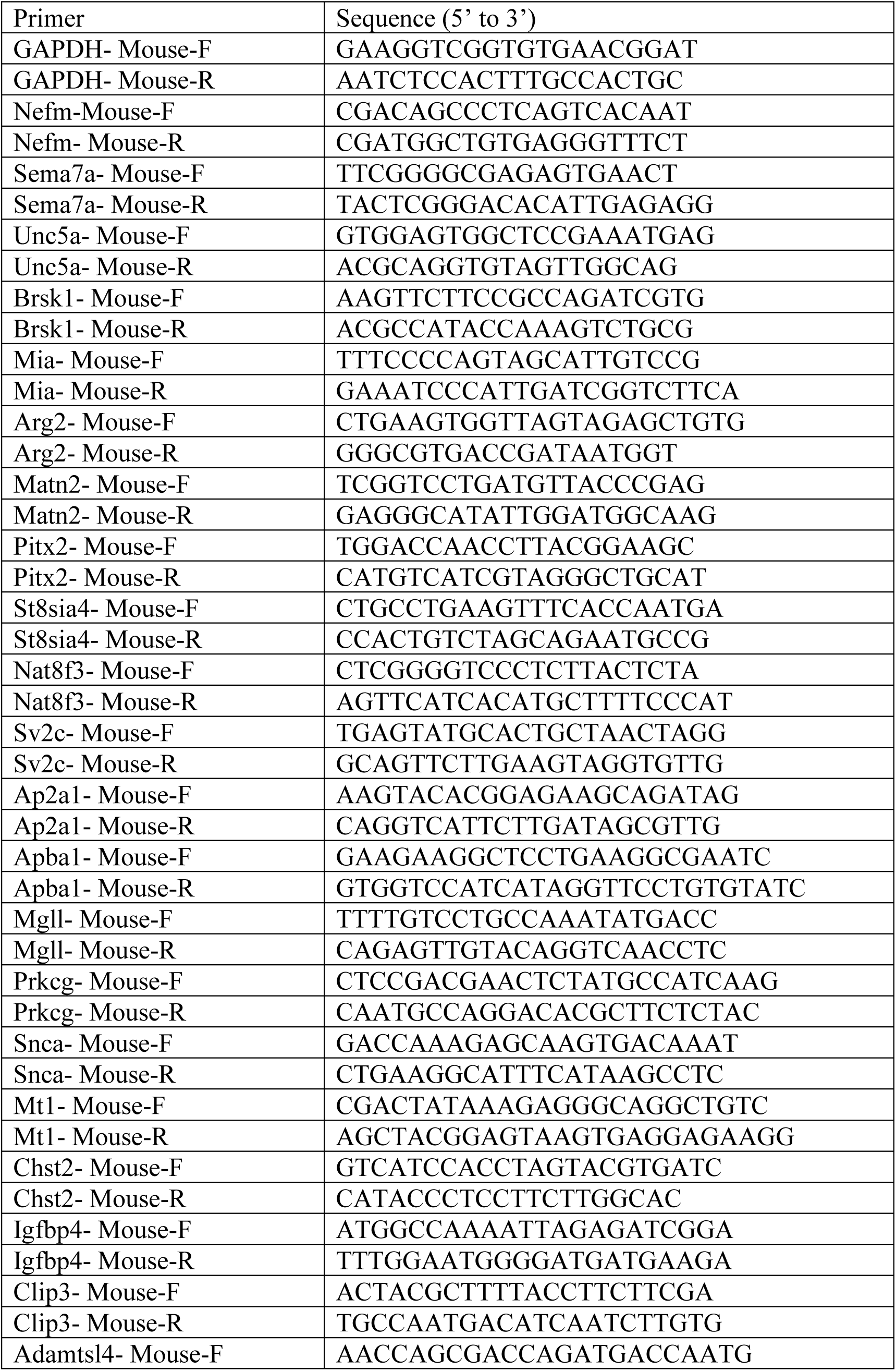
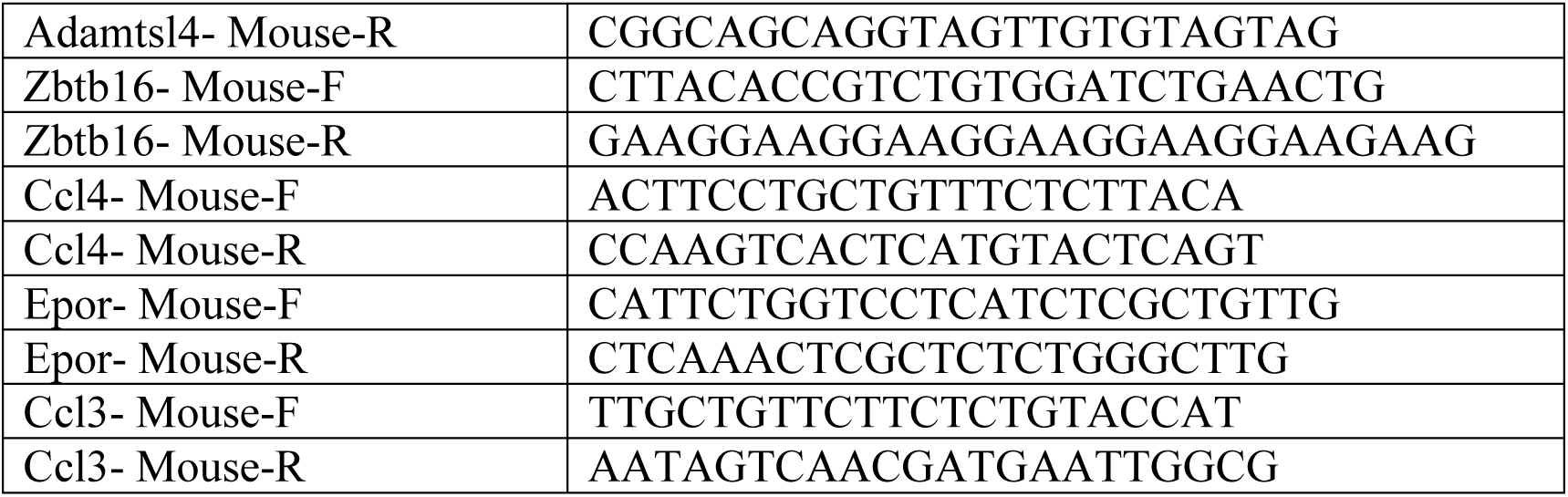
The following primers used in this study were synthesized by Sangon Biotech

### Statistical Analysis

All statistical analyses and plots were made using Prism software or MATLAB. All the computed parameters were quantified and compared between tested groups. Statistics were performed on averaged values per mouse. All data are reported as the mean ± SEM. Differences between two groups were determined with Student’s *t*-test. One-way or two-way analysis of variance followed by a post hoc Bonferroni’s or Fisher’s least significant differences (LSD) test was used for multiple comparisons. No randomization was used in our experiments. The criterion for statistical significance was *p* < 0.05.

### Data Availability Statement

The data that support the findings of this study are available from the corresponding author upon reasonable request.

## Acknowledgments

We thank Dr Zilong Qiu for the gift of *ChAT^cre^* knock-in mice. Our study was financially supported by the National Natural Sciences Foundation of China (82171376, 81971164, 81330026, 81771330), the National Key Basic Research Development Program of the Ministry of Science and Technology of China (973 Program, 2013CB945600), a project funded by the Priority Academic Program Development of Jiangsu Higher Education Institutions, and the Key Research and Development Plan of Jiangsu Province (BE2018654), Shanghai Municipal Science and Technology Major Project (No.2018SHZDZX01), ZJ Lab, and Shanghai Center for Brain Science and Brain-Inspired Technology.

## Author contributions

Conceptualization: Y.L., K.Z., and D.Y. methodology: Y.L., K.Z., D.Y., T.Z., Y.Z., Y.N., and S.W.; investigation: K.Z., D.Y., W.W., H.Z., W.Y., and M.H.; visualization: K.Z., D.Y., W.W., H.Z., and W.Y.; supervision: Y.L.; writing—original draft: K.Z., W.W., and H.Z.; writing—review and editing: Y.L., K.Z., W.W., and H.Z.

---

**Figure 2-figure supplement 1.**
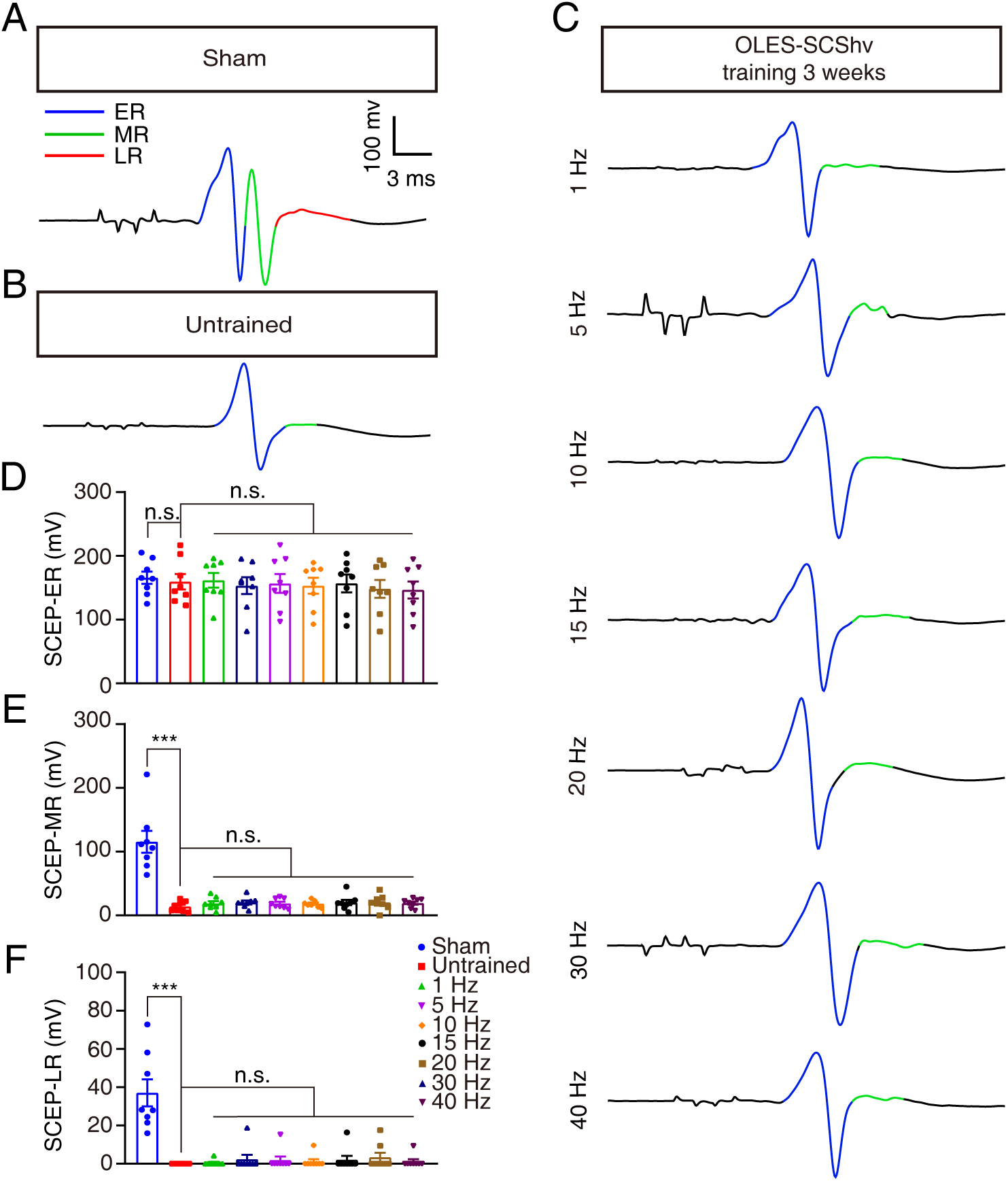
SCEPs were recorded in TA muscles after OLES- SCShv. (**A-C**) SCEPs were recorded in sham (**A**), untrained (**B**), and 1- to 40-Hz OLES-SCShv (**C**) groups. (**D-F**) The amplitudes of ERs (**D**), MRs (**E**), and LRs (**F**) of SCEPs in the 1- to 40-Hz OLES-SCShv, untrained, and sham groups (n = 8 mice). Data are shown as the mean ± SEM, ns: no statistical difference, \*\*\**p* < 0.001, one-way ANOVA followed by Bonferroni’s post-test.

**Figure 3-figure supplement 1.**
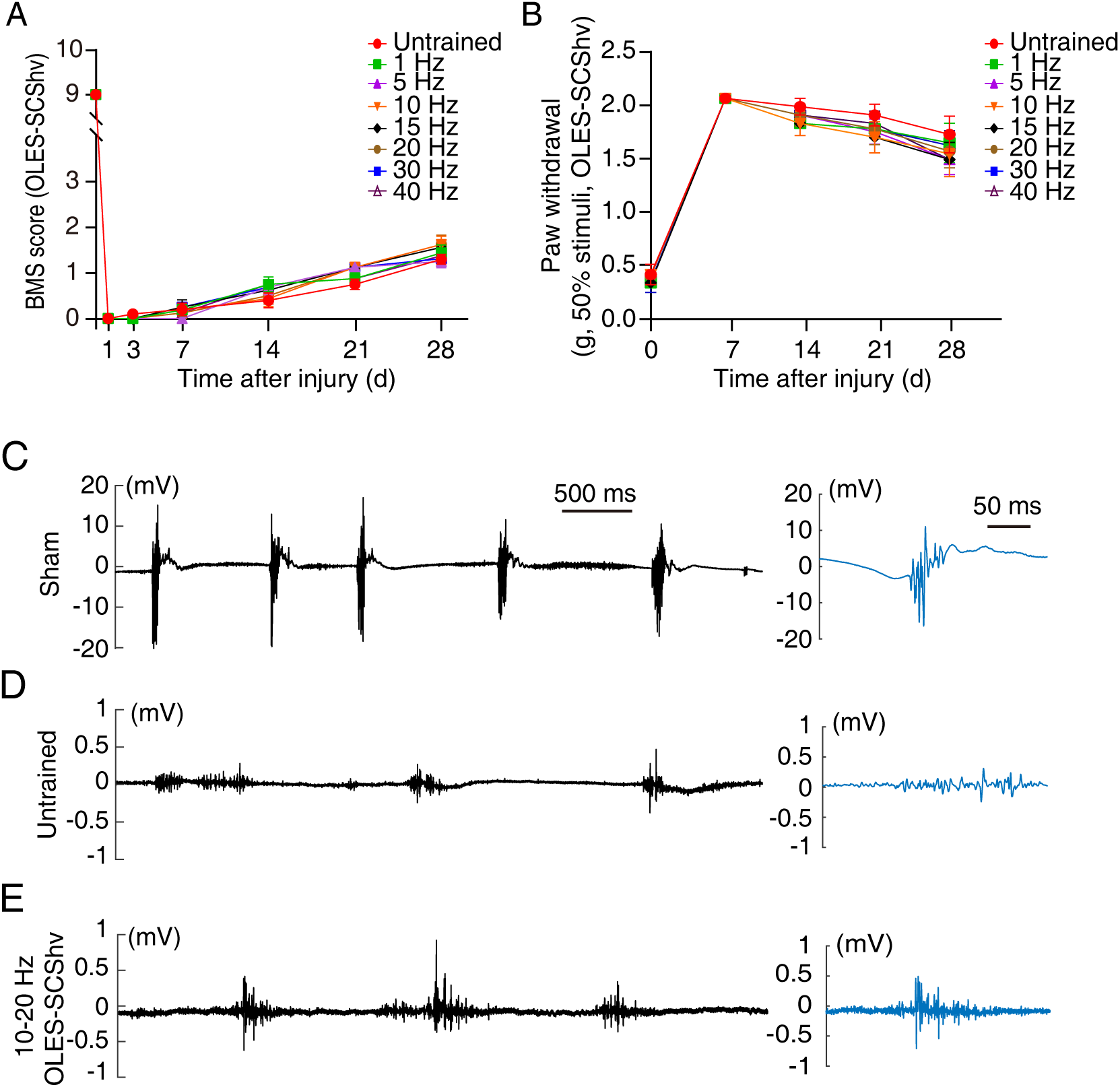
Sensorimotor function of the hind limbs in SCI mice after OLES-SCShv training. (**A**) Hindlimb BMS scores of mice in the 1- to 40-Hz OLES-SCShv groups at different time points after SCI (n = 8 mice). (**B**) The mechanical pain threshold for hind limbs in the 1- to 40-Hz OLES-SCShv groups at different time points after SCI (n = 8 mice). (**C-E**) Left, the graphs show surface EMGs for TA muscles in the sham (**C**), untrained (**D**), and 10−20 Hz OLES-SCShv (**E**) groups within 5s. Right, the representative waveform on the left. Data are shown as the mean ± SEM, two-way ANOVA followed by Fisher’s least significant differences post-test (LSD) (**A** and **B**).

**Figure 3-figure supplement 2.**
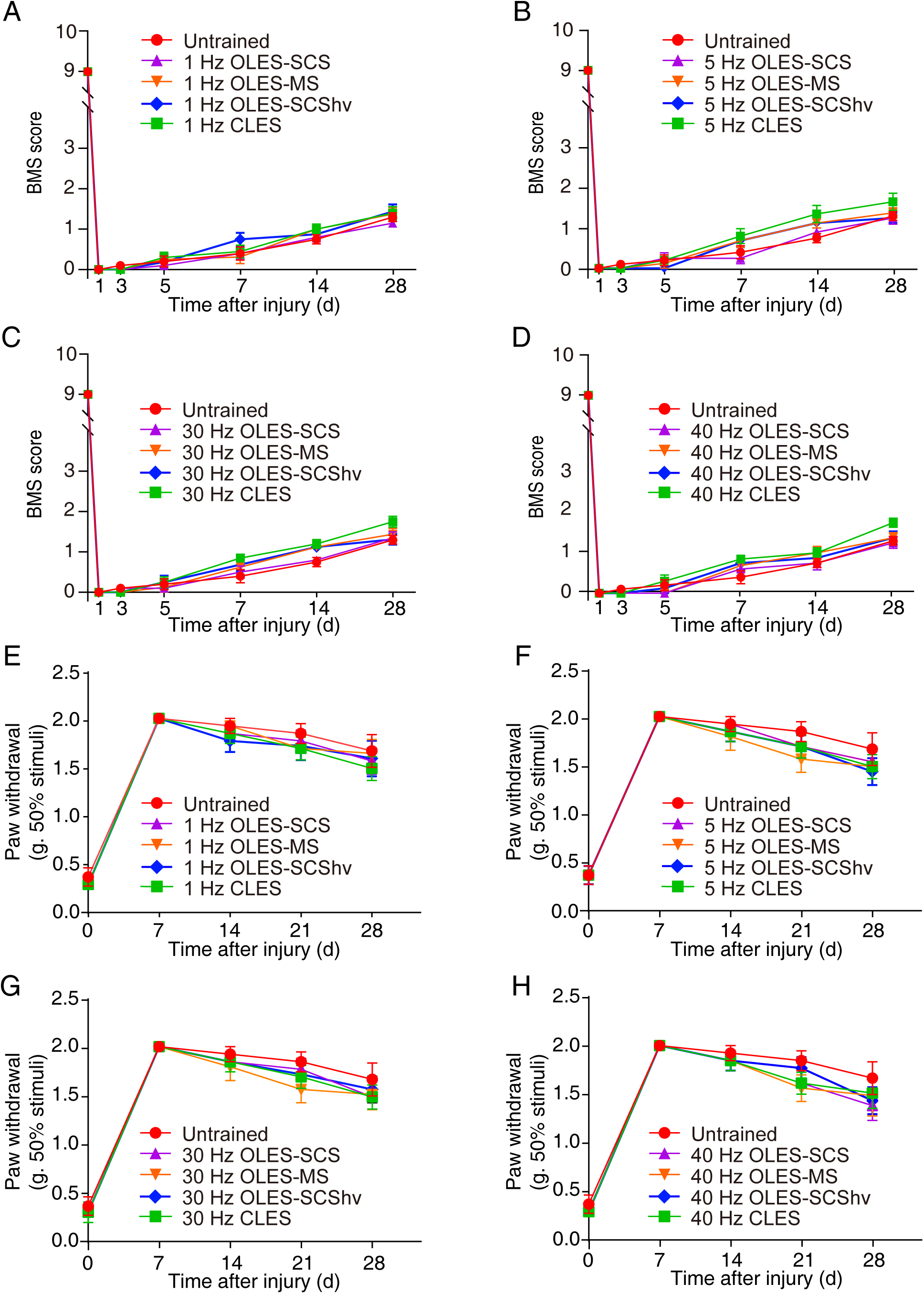
Sensorimotor function of the hind limbs in SCI mice after ineffective CLES and OLES training. (**A**-**D**) Hindlimb BMS scores of mice in the 1-Hz (**A**), 5-Hz (**B**), 30-Hz (**C**), and 40-Hz (**D**) CLES and OLES groups at different time points after SCI (n = 8 mice). (**E**-**H**) The mechanical pain threshold (as measured by paw withdrawal after application of von Frey filaments) for hind limbs in the 1-Hz (**E**), 5-Hz (**F**), 30-Hz (**G**), and 40-Hz (**H**) CLES and OLES groups at different time points after SCI (n = 8 mice). Data are shown as the mean ± SEM and two-way ANOVA followed by Fisher’s least significant differences post-test (LSD)

**Figure 4-figure supplement 1.**
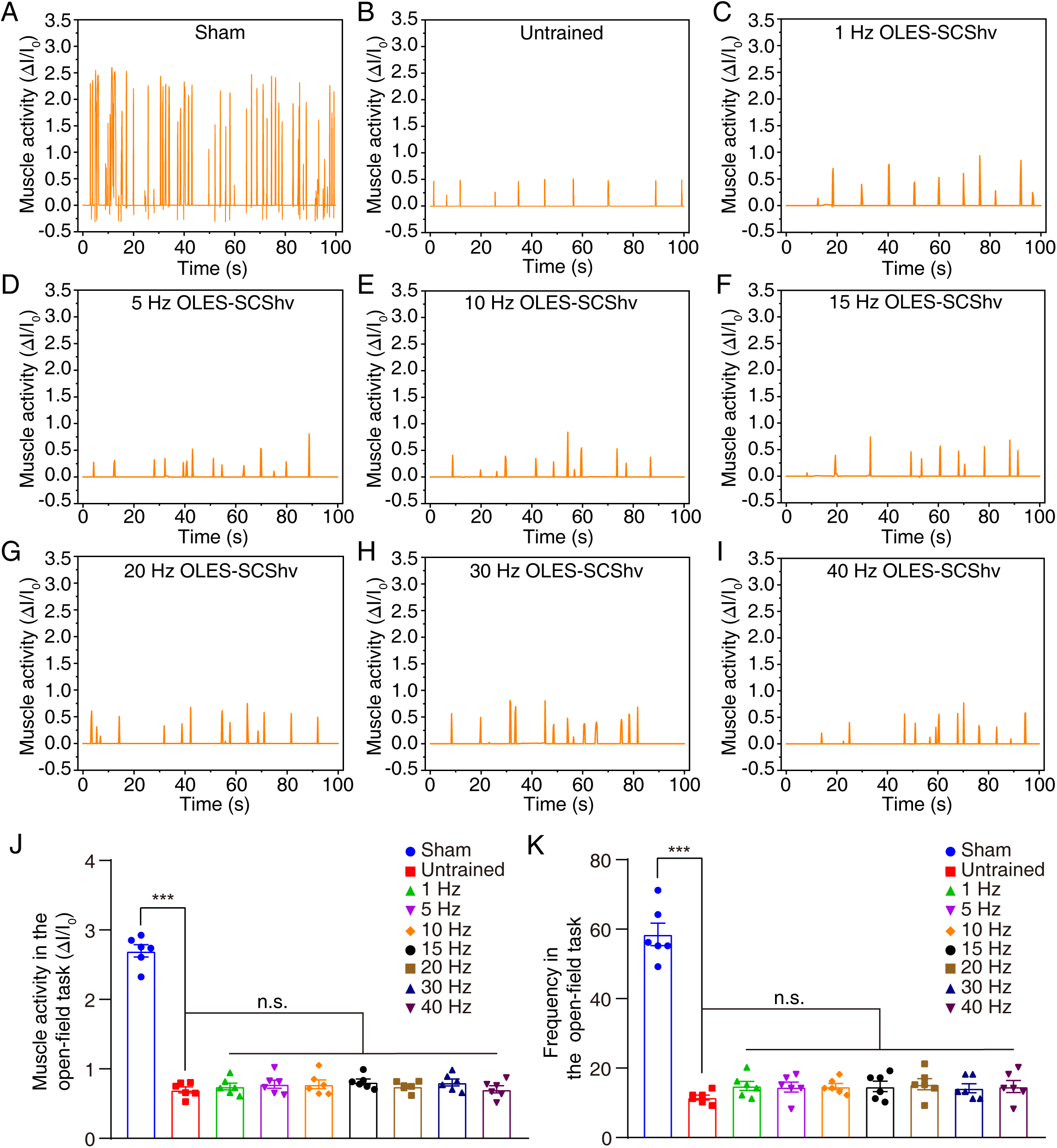
TA muscle contraction function of the hind limb in SCI mice after OLES-SCShv training. (**A**-**I**) TA muscle contraction in the hindlimb and its corresponding relative current in sham (**A**), untrained (**B**), and 1- to 40-Hz OLES-SCShv (**C**-**I**) groups. (**J** and **K**) The intensity (**J**) and frequency (**K**) of TA muscle contraction in mice in the sham, untrained, and 1- to 40-Hz OLES-SCShv groups (n = 6 mice) during the open-field task. Data are shown as the mean ± SEM, ns: no statistical difference, \*\*\**p* < 0.001, one-way ANOVA followed by Bonferroni’s post-test.

**Figure 5-figure supplement 1.**
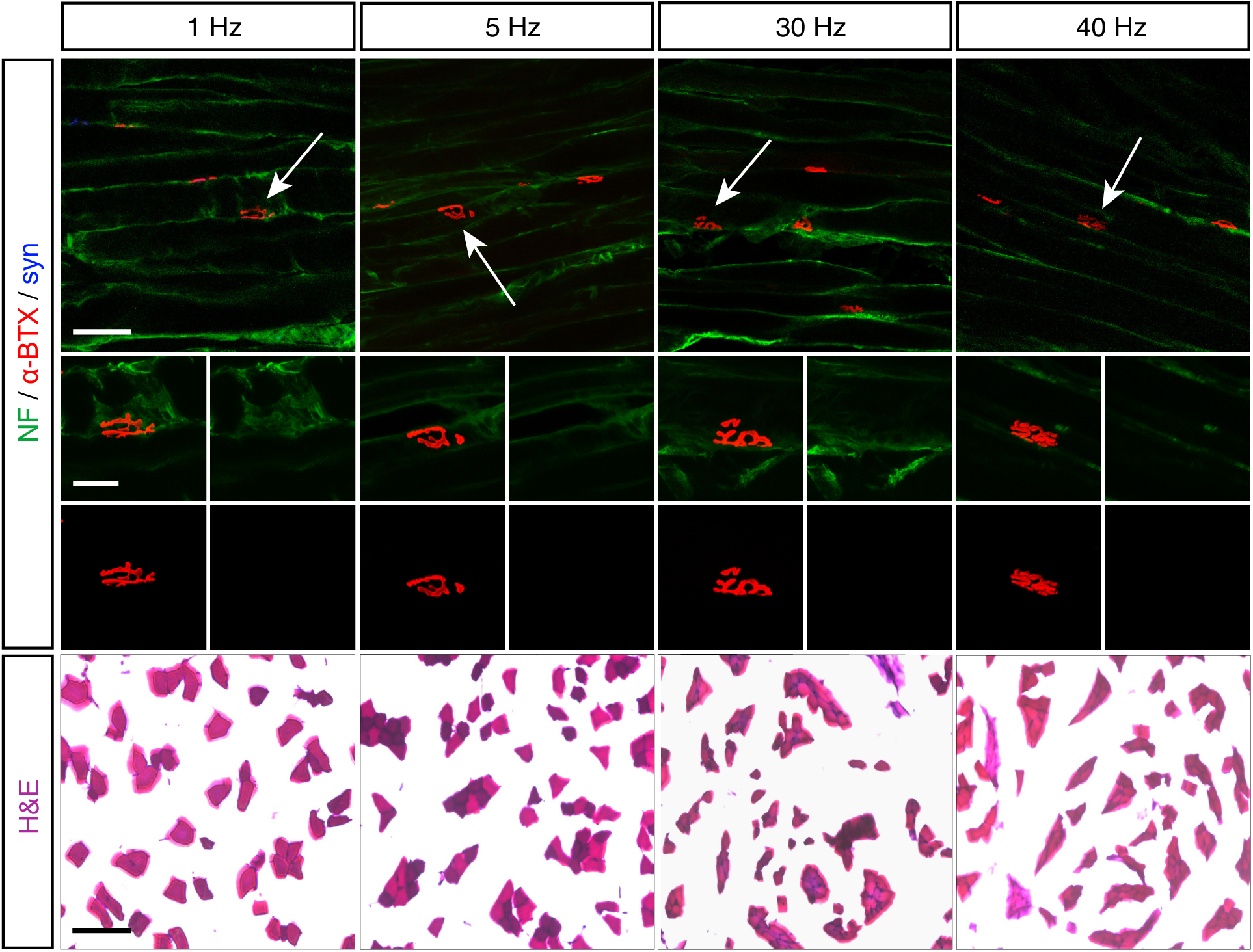
The morphology and neural innervation of TA muscle in SCI mice after ineffective CLES. Antibodies against NF (green), α-BTX (red), and synaptophysin (syn, blue) were used to label NMJs on the TA muscle of SCI mice in the 1-Hz, 5-Hz, 30-Hz, and 40-Hz CLES groups. Upper, the white arrow in the image points to a single NMJ. Scale bar, 50 μm. Middle, a higher-magnification image of the NMJ indicated by the white arrow. Scale bar, 20 μm. Lower, H&E staining of TA muscle to determine fiber cross-sectional area in the 1-Hz, 5-Hz, 30-Hz, and 40-Hz CLES groups. Scale bar, 50 μm. Quantification of these data are shown in Figure 5C and **5D**.

**Figure 6-figure supplement 1.**
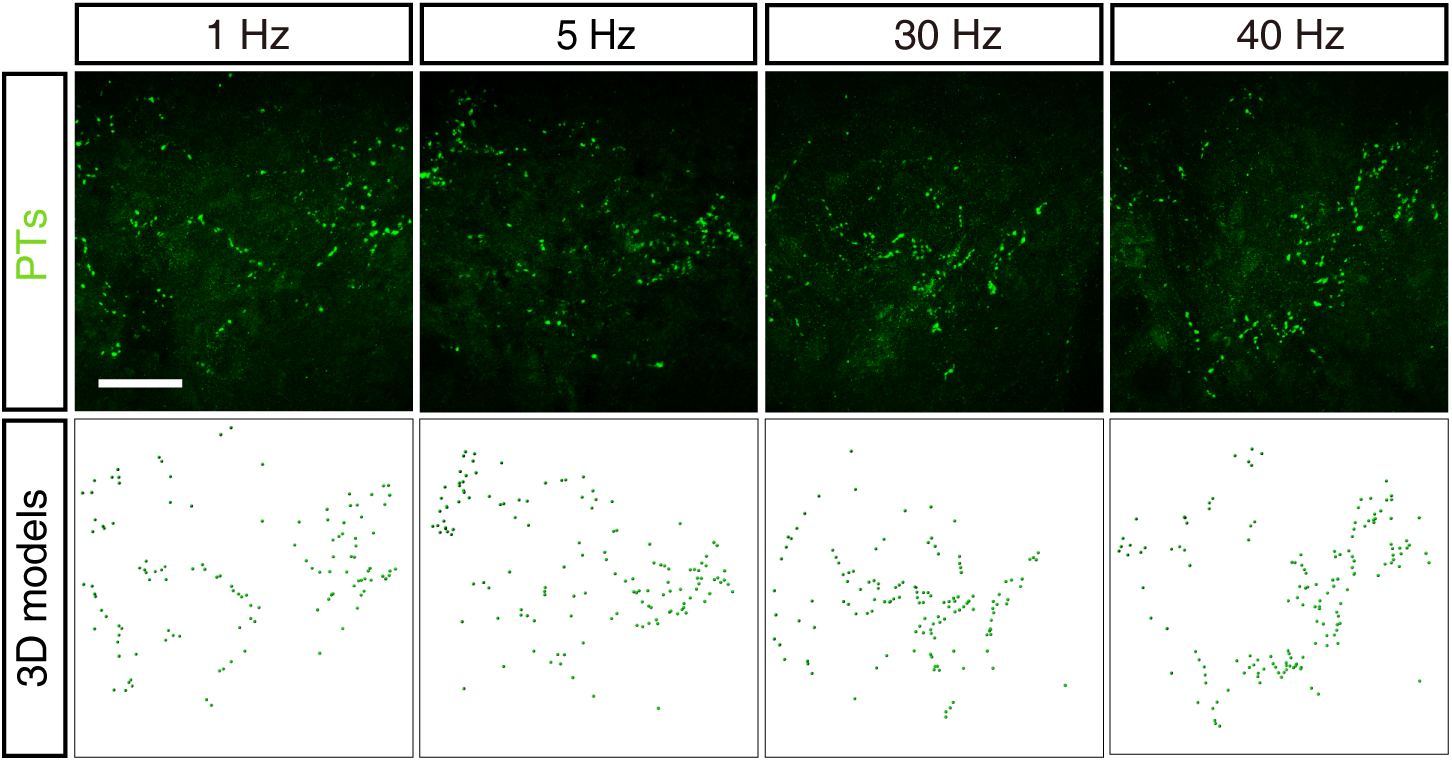
The projection of proprioceptive axons to the spinal cord in SCI mice after ineffective CLES. CTB was injected into the TA muscle of SCI mice in the 1-Hz, 5-Hz, 30-Hz, and 40-Hz CLES groups, the terminals of the proprioceptive axon were labeled (upper images), and three-dimensional modeling was performed (lower images). Scale bar, 50 μm.

**Figure 7-figure supplement 1.**
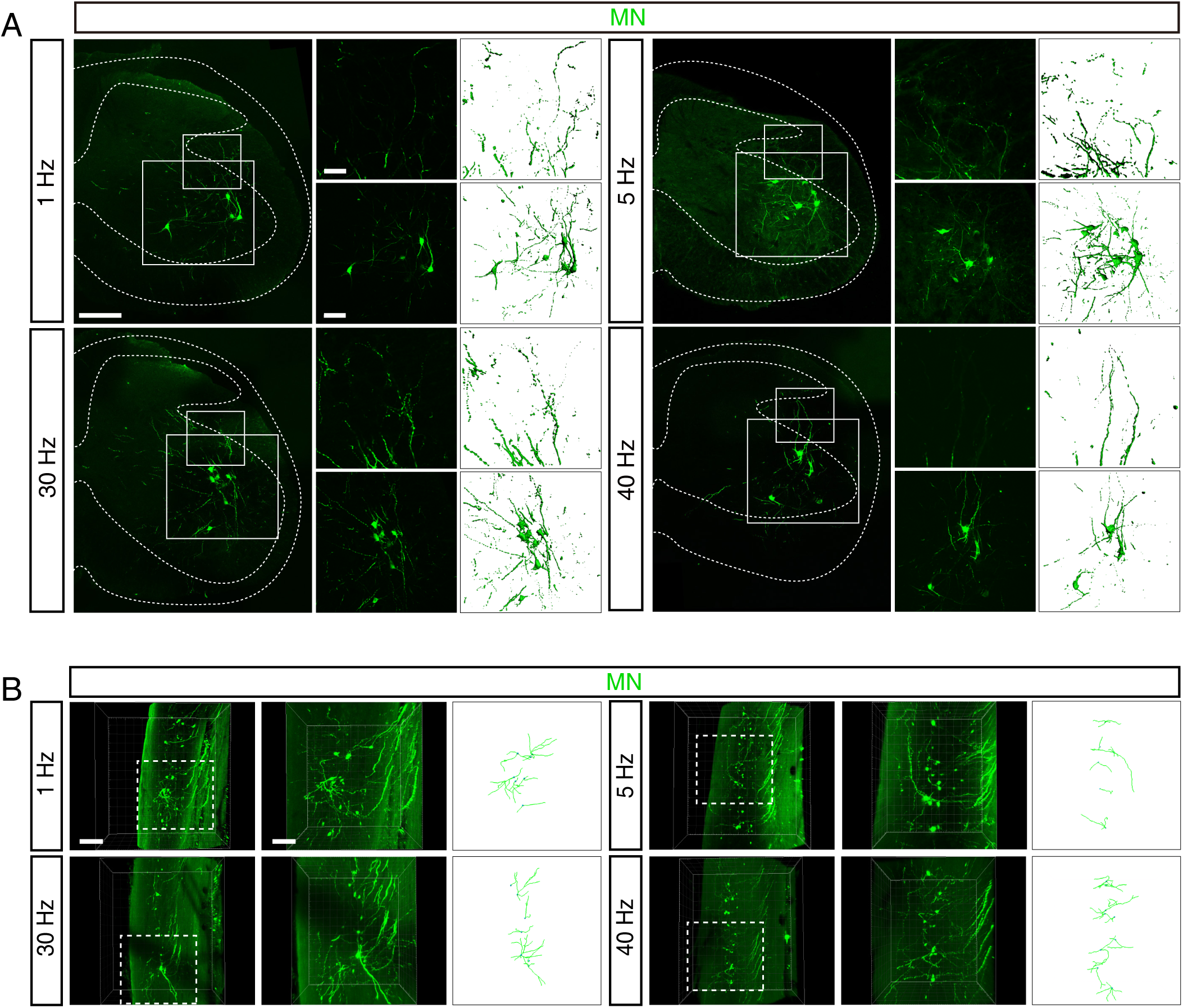
Morphological structure of motoneurons in SCI mice after ineffective CLES. (**A**) Left, spinal motoneurons of SCI mice in the 1-Hz, 5-Hz, 30-Hz, and 40-Hz CLES groups. Scale bar, 200 μm. Upper right, higher-magnification image and three-dimensional models of the dendrites in the small boxed area. Scale bar, 50 μm. Lower right, the partial image and three-dimensional model of the motoneurons in the large boxed area. Scale bar, 100 μm. (**B**) Left, spatial distribution of motoneurons of SCI mice in the 1-Hz, 5-Hz, 30-Hz, and 40-Hz CLES groups. Scale bar, 200 μm. Right, higher-magnification image and three-dimensional models of the boxed area. Scale bar, 100 μm.

**Figure 9-figure supplement 1.**
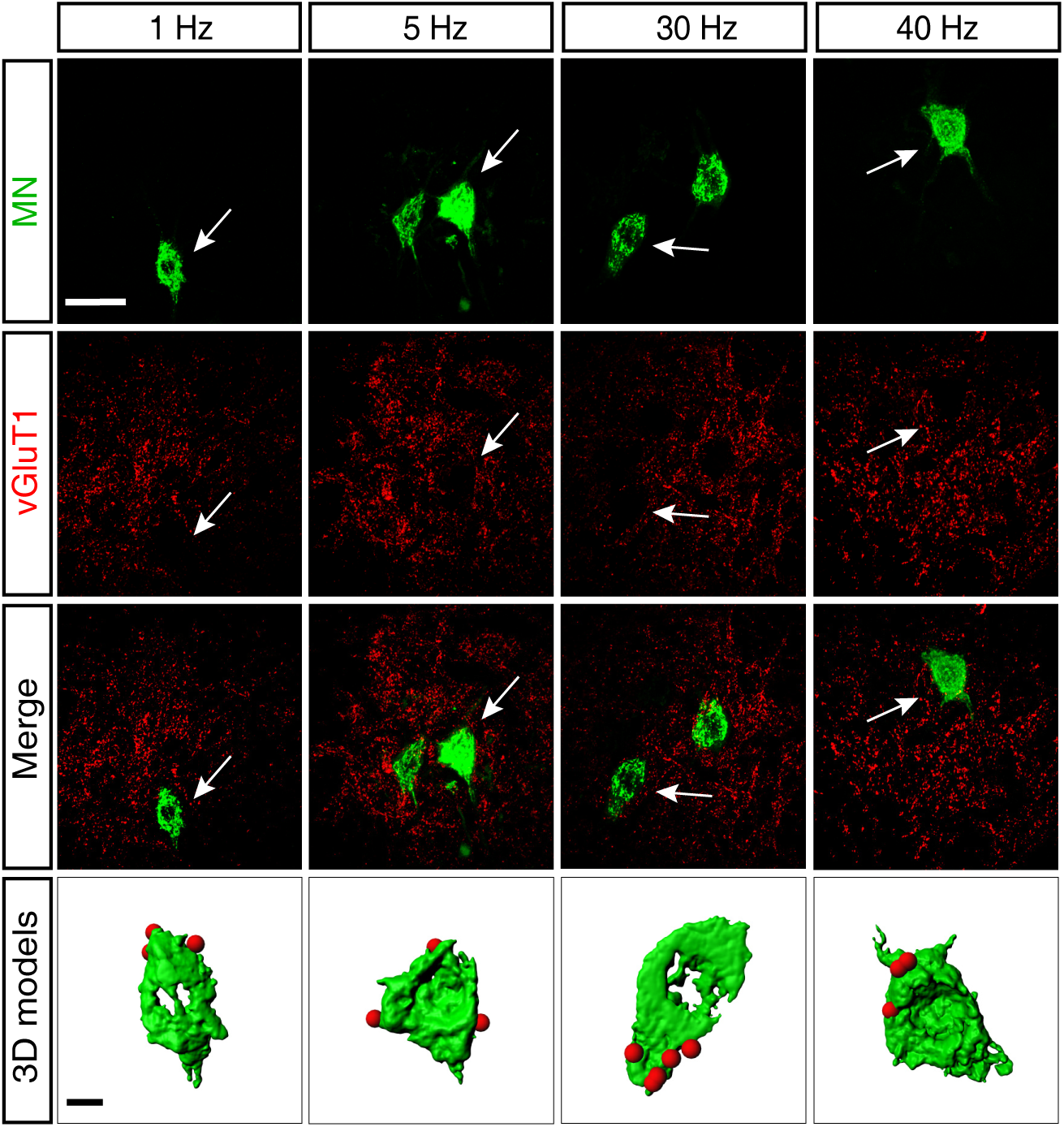
Glutamatergic synaptic input from premotor neurons to motoneurons in SCI mice after ineffective CLES. Motoneurons were marked by CTB (green) and vGluT1 (red) was used to show axonal terminals of glutamatergic neurons of SCI mice in the 1-Hz, 5-Hz, 30-Hz, and 40-Hz CLES groups. Scale bar, 50 μm. Below, three-dimensional modeling was performed on the motoneurons indicated by the white arrows and with vGluT1 on their surface. Scale bar, 10 μm.

**Figure 11-figure supplement 1.**
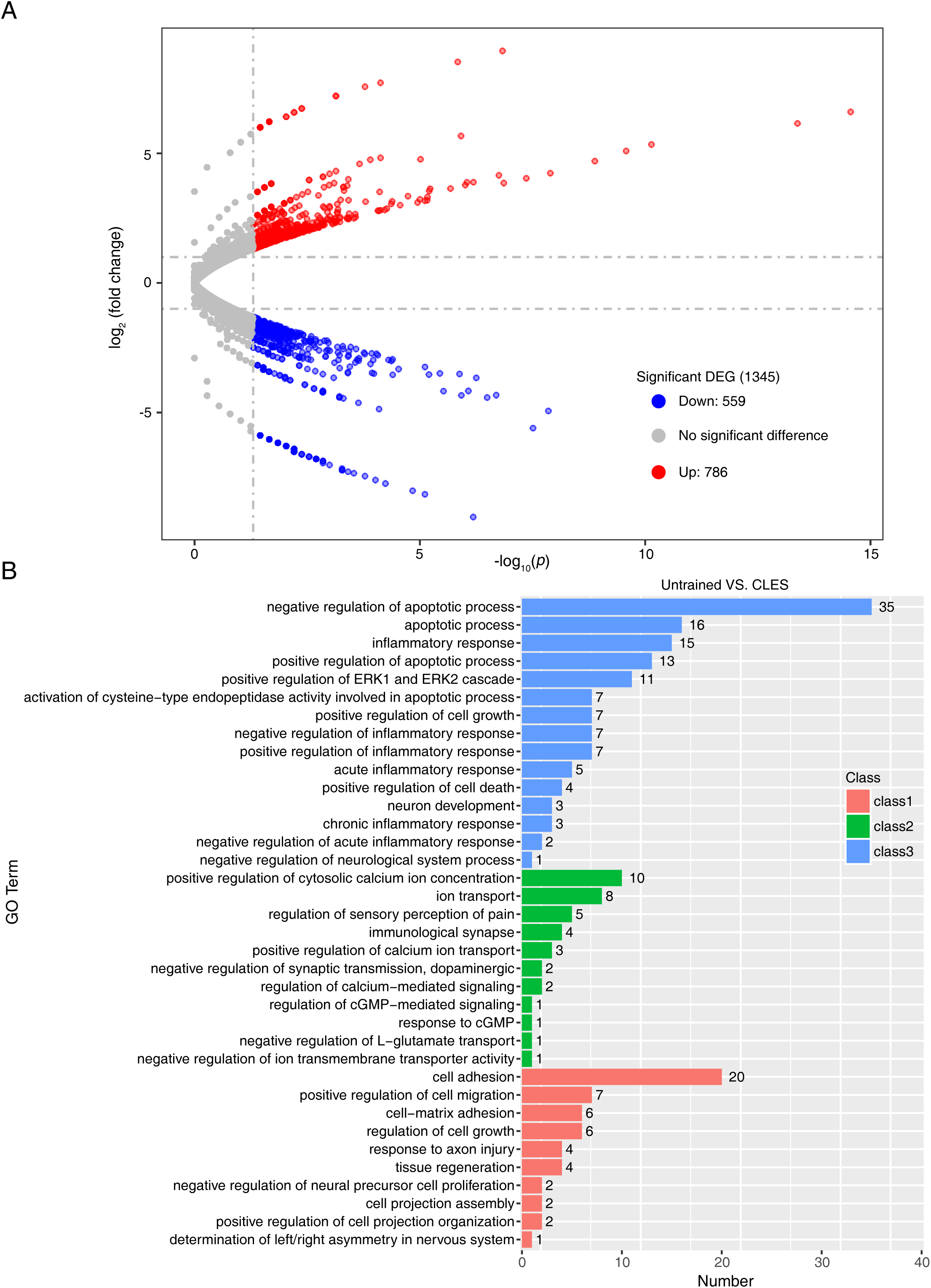
All differentially expressed genes and a partial list of gene ontology (GO) biological process terms associated with isolated spinal motoneurons. (A) All differentially expressed genes in the 10-20 Hz CLES groups as compared with the untrained group. Red dots indicate upregulated genes, gray dots indicate genes that were not significantly different, and blue dots indicate downregulated genes. (**B**) GO terms were generated for genes that were upregulated and downregulated (*p* < 0.05) in the 10-20 Hz CLES groups (The samples from the 10- Hz, 15-Hz and 20-Hz CLES groups were combined into one group) relative to their levels in the untrained group. Class 1: neuron axon development; Class 2: synaptic function; Class 3: inflammation and apoptosis.

**Figure 1- Source Data 1**

Characteristics of the CLES system.

**Figure 2- Source Data 1**

SCEPs recorded in the TA muscle of SCI mice after CLES.

**Figure 3- Source Data 1**

Detection of sensory and motor function of hind limbs in SCI mice after CLES and OLES.

**Figure 4- Source Data 1**

Evaluation of TA muscle contraction function in SCI mice after CLES and OLES.

**Figure 5- Source Data 1**

The morphology and neural innervation of TA muscles in SCI mice after CLES.

**Figure 6- Source Data 1**

Sensory-motor connectivity in SCI mice after CLES.

**Figure 7- Source Data 1**

Morphological characteristics of spinal motoneurons in SCI mice after CLES.

**Figure 8- Source Data 1**

The connection between spinal interneurons and motoneurons in SCI mice after CLES.

**Figure 9- Source Data 1**

Remodeling of glutamate synaptic connections between spinal neurons in SCI mice after CLES.

**Figure 10- Source Data 1**

Recorded Ca^2+^ signals of lumbar motoneurons in SCI mice after CLES.

**Figure 11- Source Data 1**

Changes in the transcripts of motoneurons in the spinal sensorimotor circuits of SCI mice after CLES.

**Figure 2-figure supplement 1- Source Data 1**

SCEPs were recorded in TA muscles after OLES-SCShv.

**Figure 3-figure supplement 1- Source Data 1**

Sensorimotor function of the hind limbs in SCI mice after OLES-SCShv training.

**Figure 3-figure supplement 2- Source Data 1**

Sensorimotor function of the hind limbs in SCI mice after ineffective CLES and OLES training.

**Figure 4-figure supplement 1- Source Data 1**

TA muscle contraction function of the hind limb in SCI mice after OLES-SCShv training.

**Figure 5-figure supplement 1**

The morphology and neural innervation of TA muscle in SCI mice after ineffective CLES.

**Figure 6-figure supplement 1**

The projection of proprioceptive axons to the spinal cord in SCI mice after ineffective CLES.

**Figure 7-figure supplement 1**

Morphological structure of motoneurons in SCI mice after ineffective CLES.

**Figure 9-figure supplement 1**

Glutamatergic synaptic input from premotor neurons to motoneurons in SCI mice after ineffective CLES.

**Figure 11-figure supplement 1- Source Data 1**

All differentially expressed genes and a partial list of gene ontology (GO) biological process terms associated with isolated spinal motoneurons.

